# Early steps in the evolution of vertical transmission revealed by a plant-bacterium symbiosis

**DOI:** 10.1101/522367

**Authors:** Frédéric De Meyer, Bram Danneels, Tessa Acar, Rado Rasolomampianina, Mamy Tiana Rajaonah, Vololoniaina Jeannoda, Aurelien Carlier

## Abstract

Various plant species establish intimate symbioses with bacteria within their aerial organs. The bacteria are contained within nodules or glands often present in distinctive patterns on the leaves in what is commonly referred to as leaf nodule symbiosis. We describe here a highly specific symbiosis between a wild yam species from Madagascar, *Dioscorea sansibarensis* and bacteria of the species *Orrella dioscoreae*. Using whole genome sequencing of plastid and bacteria from wild-collected samples, we show phylogenetic patterns consistent with a vertical transmission of the symbionts. Unique among leaf nodule symbioses, the bacteria can be cultured and are amenable to comparative transcriptomics and phenotypic characterization, revealing a potential role in complementing the host’s arsenal of secondary metabolites. We propose a very recent acquisition of the vertical mode of transmission in this symbiosis which, together with a large effective populations size explain the cultivability and remarkable lack of genome reductive evolution in *O. dioscoreae*. We leverage these unique features to reveal pathways and functions under positive selection in these specialized endophytes, highlighting the mechanisms enabling a permanent association in the phyllosphere.

## INTRODUCTION

Microorganisms can establish a wide range of beneficial interactions with plants, often contributing to mineral uptake, nitrogen fixation, or plant defense. Most of the mutualistic associations with bacteria are facultative and have been widely studied at the root level [1–3], with much less focus on the phyllosphere or endosphere despite recent findings that plants actively shape phyllosphere microbial communities [4, 5]. Furthermore, the molecular mechanisms enabling the establishment of these interactions are not well characterized outside of a few and usually pathogenic model systems [6].

Leaf nodule symbioses represent some of the most intimate associations between plants and bacteria. The symbionts reside in dedicated structures called leaf glands or nodules, and are transmitted between generations via seeds [7]. Most leaf nodule symbioses are found in species of the Rubiaceae (*Psychotria* and *Pavetta*) and Primulaceae (*Ardisia*) families, and their symbionts are members of the *Burkholderiaceae* family of β-proteobacteria. The association is essential for both hosts and symbionts: Candidatus Burkholderia (*Ca*. Burkholderia) species cannot be cultured outside of their host and bacteria-free *Psychotria kirkii* and *Ardisia* crenata display severe growth defects [8, 9]. This co-dependence between host and symbiont is likely the result of co-evolution over several million years, compounded by small effective population sizes and genetic drift [7]. Typical of vertically-transmitted symbiotic bacteria, *Ca*. Burkholderia leaf nodule symbionts show extensive signs of reductive genome evolution, with coding capacities ranging from 41.7 % to 67.3 % and an accumulation of pseudogenes and insertion sequences [10–12]. Despite extensive genome erosion, some symbionts have been shown to produce secondary metabolites, likely involved in the protection of the host from herbivory, such as the insecticidal kirkamide and the depsipeptide FR900359, as well as the herbicidal streptol-glucoside possibly involved in allelopathic interactions [12–14]. Because of genomic instability and evolved co-dependence, it is unclear whether secondary metabolism was present in the ancestor of leaf nodule *Burkholderia* or acquired as a secondary trait.

We recently described a leaf nodule symbiosis in the monocot species *Dioscorea sansibarensis* [15]. *D. sansibarensis*, or the Zanzibar yam, is a true yam native to Madagascar and tropical Africa [16]. This fast growing vine, like many yam species, reproduces asexually through aerial bulbils and underground tubers and is not known to produce viable seeds [17]. Leaves of *D. sansibarensis* display prominent acumens or ‘drip-tips’, harboring high titers (>10^9^ cfu/g) of *Orrella dioscoreae* (*O. dioscoreae*), a newly described species of the *Alcaligenaceae* family [15]. Similar to leaf nodule symbioses in dicot species, the bacteria are hosted extracellularly in the leaf gland and do not invade the mesophyll or vasculature of the host and do not spread systemically [7]. In contrast to other leaf nodule symbioses, *O. dioscoreae* can be cultured [15].

The aim of this study was to (i) characterize the prevalence and mode of transmission of *O. dioscoreae* in wild populations of *D. sansibarensis*; (ii) propose hypotheses regarding the recruitment of functions in leaf nodule symbiosis and (iii) leverage the unique tractability of the *D. sansibarensis* leaf symbiosis to uncover the characteristics of a strict endophytic lifestyle. We show that the association with *O. dioscoreae* is ubiquitous and highly specific in *D. sansibarensis*. We provide evidence that the symbiosis and vertical transmission evolved during the Pleistocene, offering a unique opportunity to document the early events shaping the evolution of a hereditary plant-microbe symbiosis. Finally, secondary metabolism seems to play a central role in the *D. sansibarensis* leaf nodule symbiosis, suggesting that the acquisition of novel metabolism is a pre-requisite for the evolution of symbiont capture in the phyllosphere.

## MATERIAL AND METHODS

### Bacterial strains and growth conditions

All *O. dioscoreae* strains (Table S1) were grown at 28°C on tryptic soy agar (TSA) medium or AB minimal medium [18] supplemented with 20 g/L sodium citrate and 0.5 g/L yeast extract unless otherwise indicated. Aerobic cultures were grown with vigorous shaking (200 rpm) in 500 mL Erlenmeyer flasks containing 100 ml of medium for 20h.

### Sampling and identification of wild *Dioscorea sansibarensis* samples

Leaf nodule samples from wild *Dioscorea sansibarensis* plants were collected from 12 different sites in Madagascar during two field collections in November 2016 and May 2017, with research permit 158/16/MEEF/SG/DGF/DSAP/SCB.Re issued by the Ministry of Environment, Ecology and Forests of the Republic of Madagascar. At each sampling location, about ten leaf nodules from distinct plants were harvested. Samples were immediately placed in sealed plastic sampling bags containing 5-10g of silica gel (Carl Roth) for dehydration and shipping. The GPS coordinates of the sampling locations are given in Table S2. Appropriate measures were taken to comply with Nagoya protocol guidelines.

### DNA-extraction and PCR

Silica-dried samples were processed using a combination of bead beating (Retsch MM400, Haan, Germany) and the Maxwell^®^ 16 DNA Purification Kit (Promega, Madison, WI, USA) and are described in detail in supplementary information. PCR amplification and sequencing of the *nrdA* gene was used to confirm the presence of *Orrella dioscoreae*. PCR analysis using primers specific for the chloroplastic markers *matK*, *rbcL* and *rpl32-trnL* were used to confirm plant species against reference sequences obtained from a vouchered *D. sansibarensis* specimen from the live collection of the botanical garden of Ghent University (accession 19001189). All oligos used in this study are listed in Table S3.

### Metagenome assembly and annotation

Sequencing reads were prepared for assembly by adapter trimming and read filtering using Trimmomatic [19], removing reads with phred scores below 30 and discarding non-paired reads. To assemble sequencing reads derived from *Orrella dioscoreae*, an approach based on Albertsen *et al*. [20] was used. In short, filtered reads were assembled using SPAdes v3.10.1 [21] in metagenome mode using kmer-lengths of 21, 33, 55, and 77. The read coverage and GC-content of the resulting contigs were calculated and plotted using the Matplotlib package in Python [22]. Taxonomic classification of the contigs was done using the Kraken software and overlaid on the plot [23]. Contigs consistent with *O. dioscoreae* were selected and re-assembled as previously described using SPAdes in careful mode, using kmer-lengths of 21, 33, 55, 77, 99, and 121 [12]. Assembly statistics of the resulting assemblies were generated using Quast v4.5 [24]. Contigs smaller than 500 nt, with low coverage (< 1/3 of average coverage), or classified as eukaryotic were discarded from the final assembly. Annotation was performed with the RAST online service [25] with gene prediction enabled. Orthologs were computed using OrthoMCL v1.4 [26], using a Blastp e-value cut-off of 1.0×10^−6^, 50% identity over 50% query length, and an inflation factor of 1.5. EGGNOGmapper was used to assign GO, EggNOG and COG category annotations to the proteins [27]. Analysis of putative secondary metabolite gene clusters, including NRPS adenylation domain substrate prediction were done using the AntiSMASH web server [28]. Analysis and phylogenetic clustering of NRPS condensation domains was done using the NaPDoS web server [29]. Genome comparisons were done using the NCBI blastn program and blast ring diagrams were drawn using the Circos v0.63 software [30]. Sequencing reads and genome assemblies were deposited in the European Nucleotide Archive under accession PRJEB30075.

### Microbial diversity analysis of leaf nodules

To assess the diversity of bacteria within the leaf nodule, the WGS reads of each nodule were classified using Kraken [23], using a custom kraken database built from the ‘bacteria’ and ‘plastid’ components of the NCBI RefSeq database (downloaded Feb 2017). Additionally, sequencing reads were analysed using MetaPhlAn2 [31] for detection of microbial eukaryotes. Strain-level diversity of *O. dioscoreae* within one plant was assessed by WGS of 20 nodules sampled from a single *D. sansibarensis* specimen kept in the greenhouse of the botanical garden of Ghent University. Total DNA was extracted, pooled in 4 pools of 5 samples and sequenced using shotgun methods as described above. The resulting WGS reads were trimmed and filtered as described above and mapped to the repeat-masked reference genome sequence. Reads mapping to multiple sites in the genome were discarded from the analysis. Polymorphic sites were detected using CLC genomics workbench v7.5, using a base quality filter of 25, with a minimum quality score of 20 in the neighbourhood of 5 bases. Only polymorphisms supported by at least 5 reads were considered.

### Gnotobiotic culture of *D. sansibarensis*

Five *D. sansibarensis* bulbils were collected from the greenhouse of the botanical garden of Ghent University, thoroughly washed with tap water, surface sterilized with 70% ethanol and 1.4% sodium hypochloride for 5 min each, and rinsed three times with sterile MilliQ water. Each bulbil was transferred to an autoclaved (121°C for 15min) microbox container (Combiness, Belgium) containing half-strength Murashige and Skoog (MS) medium and incubated for 2 months at 28°C with a 16h photoperiod and light intensity of 50 μmol m^−2^ s^−1^. After germination, e leaf nodules were aseptically dissected and ground in sterile 0.4% NaCl. The macerate was streaked on TSA medium and incubated at 28°C for 48h. Identification and typing of isolates was done by colony PCR amplification and sequencing of the *nrdA* gene as described above.

### RNA isolation and sequencing

*Dioscorea sansibarensis* were grown in the greenhouse at 25°C with a 14h-light/10h-dark-photoperiod cycle (approx. 75 μmol m^−2^ s^−1^ at time of harvest). Nodules from 3 three-month-old plants were dissected, at the middle of the light phase (2 pm) and middle of dark phase (1 am), using sterile scissors decontaminated with RNAase ZAP (Sigma Aldrich, St. Louis, MI, USA). Each pair of collected nodules (day/night), were collected from the same plant. RNA was isolated using the Aurum^™^ Total RNA Mini Kit (BioRad, USA) according to manufacturer’s recommendations. Stranded cDNA libraries were constructed and sequenced at the Wellcome Trust Centre for Human Genetics (Oxford, UK) (see Supplementary information for detailed methods). Analysis of the resulting sequencing reads was done using the DEseq2 software using the LMG 29303^⊤^ reference genome [32].

### Phenotypic analysis

*O. dioscoreae* was grown on R2A agar (Oxoid) and grown overnight at 28°C. Cells were suspended in the inoculation fluid IF-0 (Biolog, Hayward, CA) supplemented with dye A to a final turbidity of 85%T according to manufacturer’s recommendation. The suspension was then inoculated on Biolog plates PM1 and PM2A and incubated at 28°C for 48 h under aerobic conditions. Development of colour indicating substrate respiration was monitored at 24h and 48h.

## RESULTS

### The *D. sansibarensis* / *O. dioscoreae* association is prevalent in nature

We previously reported the isolation of *O. dioscoreae* from several *D. sansibarensis* specimens from European botanical gardens [15]. To investigate whether the association is also prevalent in nature, we isolated total DNA from 47 *D. sansibarensis* leaf acumens collected in 12 sites in the eastern Atsinanana and northern Diana regions of Madagascar (Figure 1). We could detect the presence of *O. dioscoreae* DNA by PCR in all samples using primers specific to the *nrdA* gene sequence of *O. dioscoreae* LMG 29303^⊤^. Sequence analysis of the *nrdA* PCR products revealed very low diversity among samples, with a minimum of 97.6 % identity to the type strain LMG 29303^⊤^. To investigate the prevalence of *O. dioscoreae* inside the leaf nodule, we generated shotgun metagenome sequences for 20 samples representative of the sites sampled and *nrdA* sequence types (Figure S1). *O. dioscoreae* was the only microbial species consistently found in all samples (Figure S2), comprising on average 95.65% of reads classified as bacterial (min 92%; max 98%). Furthermore, *O. dioscoreae* has never been isolated outside of *D. sansibarensis* leaf nodules and an exhaustive search for *Orrella* rRNA sequences in 1677 samples from 71 public metagenome projects failed to retrieve sequences above the 98% identity threshold (Supplementary information). A Blast search of the NCBI nr and rRNA databases using the 16S sequence of *O. dioscoreae* LMG 29303^⊤^ yielded four hits with >99% nucleotide identity. All corresponded to uncultured sequences found in the midgut of the cicada *Meimuna mongolica* [33]. Cicadas are phytophagous sap-sucking insects, and there is a distinct possibility that *Orrella* bacteria were accidentally ingested upon feeding.These results suggest that *O. dioscoreae* is limited to its unique niche in *D. sansibarensis* and that the association is prevalent and specific in nature.

**Figure 1.**
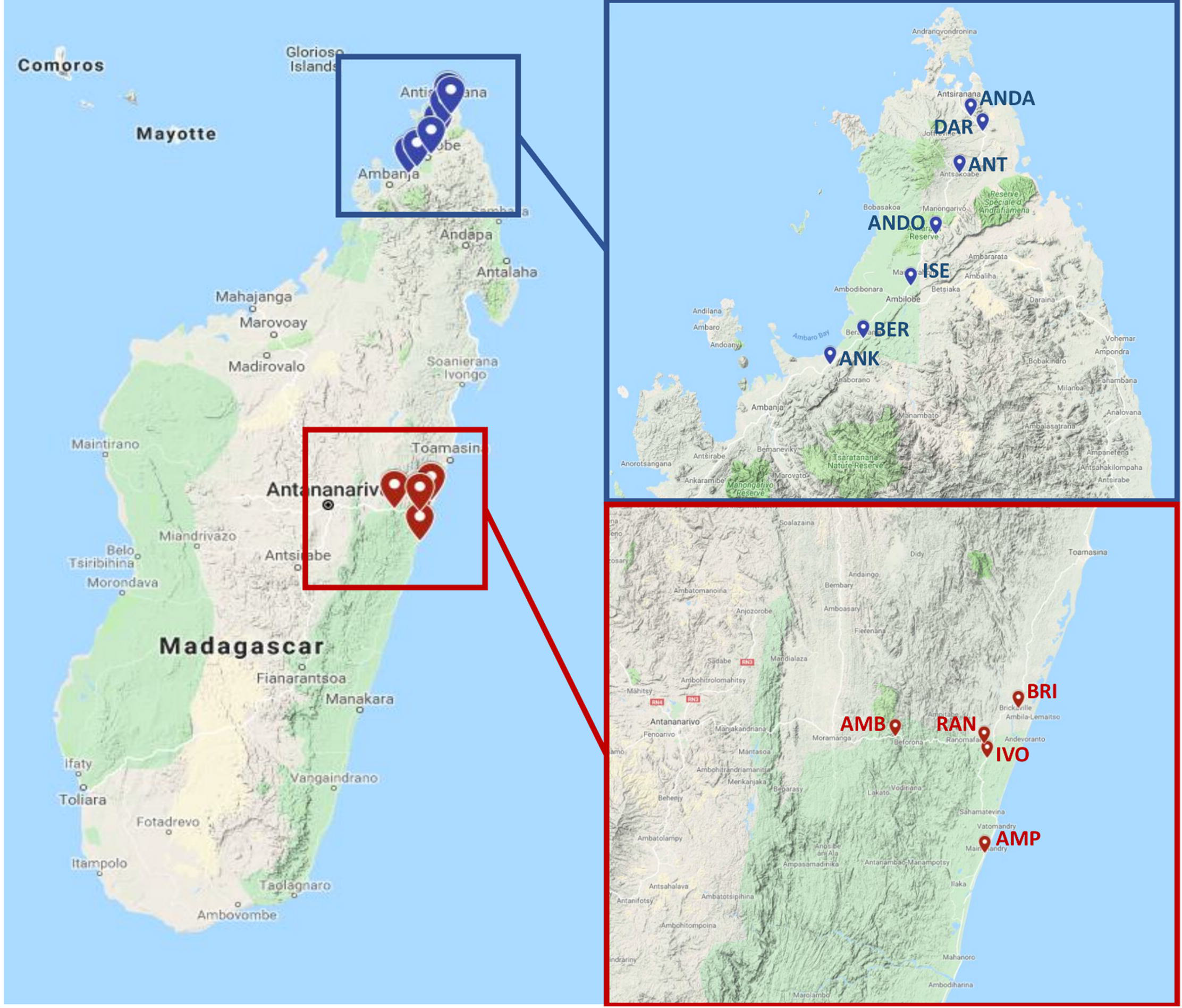
Sample site locations of collected *D. sansibarensis* leaf nodules. Red markers represent collection sites in the Atsinanana region accessed in November 2016. Blue markers represent sites in the Diana region accessed in May 2017.

### Within-host population structure of *O. dioscoreae*

Populations of *O. dioscoreae* can reach up to 10^11^ symbiont cells within a single host [15] and have the potential to display high levels of genetic diversity due to *de novo* mutations or mixed infections. To investigate the diversity of *O. dioscoreae* in one host plant, we sequenced the contents of 20 nodules from a single host, achieving an average coverage of the *O. dioscoreae* reference genome in excess of 600x. In total, we found 216 SNPs and 67 insertion-deletions. Of these, only 5 SNPs were represented in more than 10% of the reads mapped at the site. Intra-host diversity is thus very low, with low frequencies of individual SNPs indicating that *de novo* mutational processes drive intra-host diversity rather than co-infection by multiple strains.

### Dominant vertical mode of transmission of *O. dioscoreae*

Low intra-host diversity in symbiont populations may be the result of strict controls on infection or a vertical mode of transmission [34, 35]. The lack of evidence for an environmental reservoir of *O. dioscoreae* also suggests that *O. dioscoreae* cannot easily be acquired horizontally from the environment. To test if *O. dioscoreae* is transmitted vertically, germinated surface sterilized bulbils under gnotobiotic conditions. The bulbils all gave rise to plants colonized by bacteria with identical *nrdA* gene sequences to the parent (data not shown). Furthermore, *O. dioscoreae* could be isolated from macerated, surface-sterilized bulbils in high numbers (on average 2.2 × 10^5^ ± 1.2 × 10^5^ cfu/g or about 4.5 × 10^5^ per bulbil), as well as from axillary buds from which bulbils emerge. Furthermore, co-phylogenetic analysis of chloroplast and symbiont genomes of wild-collected samples revealed broad patterns of co-speciation, with a distinct biogeographical component. First, chloroplast whole genome phylogenetic analysis resolved two distinct clusters according to sampling location in the Atsinanana and Diana regions (Figure 2B). Symbiont whole genome phylogenies displayed partial congruence with the chloroplast phylogeny and a general conservation of the biogeographic signal (Figure 2A). However, statistical analysis rejected strict co-speciation between host and symbionts (P > 0.9), while reconciliation analysis introduced 4 co-speciation events, 2 losses and 3 host-switching events (Figure 2B). These data are consistent with a dominant vertical mode of transmission with occasional horizontal or host-switching events.

**Figure 2.**
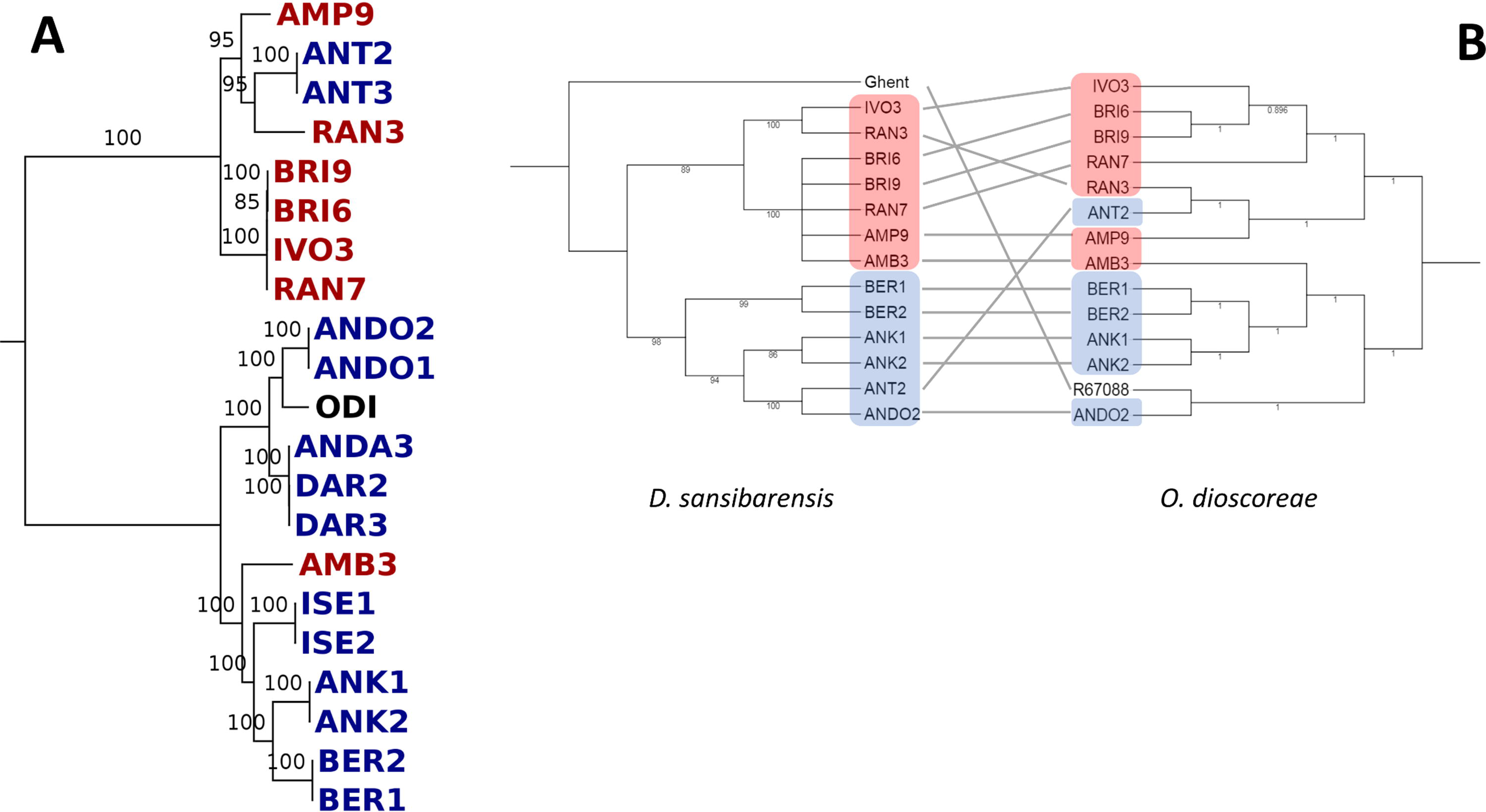
Population structure of the *D. sansibarensis*/*O. dioscoreae* symbiosis. A) Bootstrapped phylogeny of *O. dioscoreae* samples based on single copy core genes. Highlighted in red: samples collected in the Atsinanana region; highlighted in blue: samples collected in the Diana region; Black: Typestrain LMG29303^⊤^. B) Co-phylogenetic analysis of *D. sansibarensis* (left) and *O. dioscoreae* (right). Host phylogeny was reconstructed based on a concatenated SNP-based alignment of whole chloroplast sequences including invariant sites (see supplementary information for details). Bootstrap values are shown. Nodes supported by less than 75% bootstrap values were collapsed.

### Vertical transmission without genome reduction

Reductive genome evolution, a process by which the size and coding capacity of genomes tends to shrink over time, is a nearly universal phenomenon among vertically transmitted symbionts and obligate pathogens, including leaf nodule symbionts of Rubiaceae and Primulaceae [10–12, 36, 37]. The genome of *O. dioscoreae* LMG 29303^⊤^ is of average size for the family *Alcaligenaceae* (Figure S3). Reduced genomes also commonly have lower %G+C compared to free-living relatives, but the average %G+C of 66.15% does not deviate significantly from the average of 65.34% calculated from genomes of neighbouring *Bordetella* and *Achromobacter* genomes. Coding density is also high (90%), with only 40 predicted pseudogenes and 20 putative IS elements present in the genome of the type strain LMG 29303^⊤^. This lack of evidence for genome reduction could be explained by two, non-mutually exclusive reasons: (i), the *Dioscorea* leaf nodule symbiosis evolved only recently, leaving little time for the effects of genome erosion to accrue, or (ii), a large effective population size and efficient selection accounts for the maintenance of genomic structure. To obtain a measure of the efficiency of selection, we calculated genome-wide non-synonymous to synonymous substitutions rates (*d*_*N*_/*d*_*S*_) of *O. dioscoreae*. The genome-wide *d*_*N*_/*d*_*S*_ of *O. dioscoreae* is low (*d*_*N*_/*d*_*S*_ = 0.0597) and in the reported range for free-living bacteria [38]. Furthermore, core genes do not display significantly elevated *d*_*N*_/*d*_*S*_ compared to free-living *Alcaligenaceae* species (Figure S4). These data indicate that *O. dioscoreae* experience low levels of genetic drift, typical for bacteria with large effective populations but highly unusual for vertically-transmitted symbionts [39, 40].

### Recent capture of a vertically-transmitted endophyte

A recent evolution of a vertical mode of transmission may also contribute to the lack of evidence for genome erosion. The ancestor of *D. sansibarensis* diverged from sister Malagasy yam species around 22.6 Mya [41]. However, the estimated time of divergence between *D. sansibarensis* and non-nodulated sister yam species is likely a gross overestimate of the age of the symbiosis because of incomplete sampling of extant and extinct species. Using chloroplastic whole genome sequences and calibration data from Viruel *et al*. [41], we estimated that the specimens included in our study diverged during the Pleistocene and perhaps as recently as 20 000 years ago (Supplementary information). To provide another independent estimate of the divergence time of our samples, we calculated the mutation rate of *O. dioscoreae* in a single lineage sampled at 2 years interval. Using an estimated mutation rate of 2.06 × 10^−7^ substitutions/site/year, we inferred that all *O. dioscoreae* strains (including from samples collected in continental Africa) diverged from a common ancestor about 124 000 years ago (supplementary information). This estimate falls within the confidence interval derived from the chloroplast whole genome phylogenetic analysis and supports a very recent emergence of the symbiosis.

### Extensive metabolic reprogramming during symbiosis

To elucidate the function and the nature of the metabolic exchange between host and symbiont, we generated transcriptomic data of O. dioscoreae in the leaf gland of *D. sansibarensis* compared to *O. dioscoreae* LMG 29303^⊤^ grown on minimal media supplemented with citrate and ammonia as sole carbon and nitrogen sources. A total of 1639 out of 4363 (37.5%) genes were differentially expressed (p-value < 0.05; absolute log_2_ fold change ≥ 1.5) between growth *in planta* or on minimal medium, with a balanced proportion of upregulated (834) and downregulated (805) genes across all major functional categories (Figure 3). A majority of genes belonging to COG category J (translation, ribosomal structure and biogenesis) were upregulated in the leaf nodule, while most differentially expressed genes of COG category P (inorganic ion transport and metabolism), I (lipid transport and metabolism) and K (transcription) were down-regulated, perhaps indicative of a low diversity of substrates available for growth. Of the 70 upregulated genes linked to ribosome structure and function, 17 were among the top 100 upregulated genes (> 13-fold). This higher expression of ribosome components was concomitant with an upregulation of the translation elongation factors *efp*, *tsf*, *tuf* and *fusA*, as well as more than 30% of genes belonging to COG category D involved in cell cycle control and chromosome partitioning and an upregulation of genes coding for subunits of the RNA polymerase (ODI_R0061, ODI_R0131-2). Increased synthesis of ribosomes and components of the translation, transcription and replication machinery are indicative of faster growth rates *in planta* [42].

**Figure 3.**
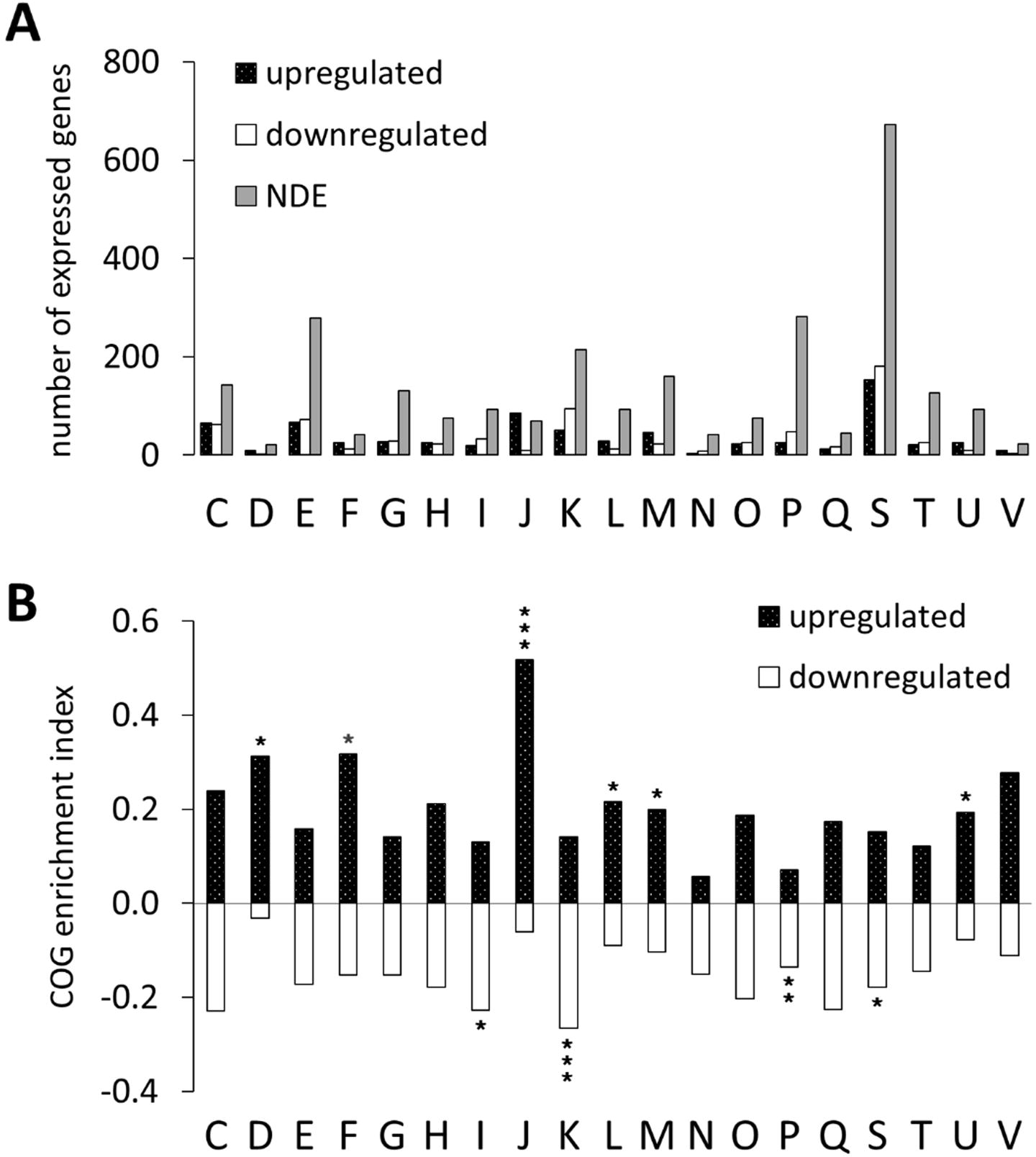
Functional profile of differentially expressed genes in the leaf nodule compared vs. culture of *O. dioscoreae*. A: Distribution of DEGs into functional COG categories. Black and white bars represent counts of up- and down-regulated genes, respectively. Grey bars indicate the total number of non-DEGs. B: COG enrichment for differentially regulated genes. The COG enrichment index is calculated by dividing the percentage of genes up-or down regulated for each category by the percentage of genes in the genome belonging to the same category. Positive values represent upregulated genes in the *in planta* growth condition (when compared with axenic growth); negative values, to the downregulated genes. Enrichment index values that are significantly different from expected (two-tailed Fisher’s exact test) are indicated by asterisks as follows: *, P≤0.05; **, P≤0.01; ***, P≤0.001. (C) = energy production and conversion; (D) = cell cycle control and mitosis; (E) = amino acid metabolism and transport; (F) = nucleotide metabolism and transport; (G) = carbohydrate metabolism and transport; (H) = coenzyme metabolism; (I) = lipid metabolism; (J) = translation; (K) = transcription; (L) = replication and repair; (M) = cell wall/membrane/envelop biogenesis; (N) = cell motility; (O) = post-translational modification, protein turnover, chaperone functions; (P) = inorganic ion transport and metabolism; (Q) = secondary structure; (S) = function unknown; (T) = signal transduction; (U) = intracellular trafficking and secretion; (V) = defence mechanisms.

### The leaf gland niche is characterized by micro-oxia, iron limitation and a sessile lifestyle

Extensive metabolic reprogramming and faster growth rates suggest that *O. dioscoreae* is highly adapted to conditions within the leaf gland environment. Upregulation of a cytochrome d ubiquinol oxidase in the leaf gland (ODI_00363-4) is reminiscent of other *Burkholderiales* bacteria grown under micro-oxic conditions and may be a response to low oxygen concentration in the leaf gland [43]. Direct free oxygen measurements taken at *Ca*. 1 mm depth beneath the leaf gland surface confirm conditions of micro-oxia, with dissolved O_2_ concentration of 35.7 ± 10.8 μM at 21°C (data not shown). Most genes related to motility and chemotaxis (ODI_R2117 - R2164) are downregulated, while genes coding for cellulose synthase (ODI_R2609-2610) and capsular polysaccharides (ODI_R0994-1001) are upregulated, all of which typical of a biofilm mode of growth [44]. Furthermore, upregulation of *fur* by more than 6-fold on average in the leaf nodule, together with upregulation of genes related to siderophore biosynthesis and uptake (ODI_R2471-78 and R2482) indicate that iron may be a factor limiting growth in the leaf gland [45–48].

### Organic acids fuel the growth of *O. dioscoreae in planta*

The genome of *O. dioscoreae* does not encode enzymes active against complex carbohydrates, and *O. dioscoreae* cannot utilize monosaccharides in culture [15]. Upregulation of the genes of the TCA in the leaf nodule by an average of 3.9-fold and a lack of overall differential expression of the pentose phosphate pathway, glycolysis or branched-chain amino acid degradation pathways (Table S4) suggest that growth inside the leaf nodule is fueled by short-chain amino acids or organic acids. To validate this interpretation, we tested growth of *O. dioscoreae* on 190 distinct carbon sources. Strain LMG 29303^⊤^ could only utilize organic acids, predominantly substrates of the TCA (Table S5). In addition, growth was supported by L-proline and L-glutamate. Two putative NADP-dependent glutamate dehydrogenases were upregulated in the leaf gland (ODI_R4072, 3.6-fold and ODI_R2231, 6.3-fold), suggesting that deamination of glutamate to 2-oxo-glutarate could provide substrates for the TCA cycle. However, a putA homolog (ODI_R1770), encoding a bi-functional proline dehydrogenase and delta-1-pyrroline-5-carboxylate dehydrogenase was not differentially regulated. This suggests that glutamate, rather than proline, possibly serves as an energy source in the leaf nodule. Growth could also be supported by D-galactonic acid (supplementary information). The *dgo* operon (ODI_R1138-1141) of D-galactonate utilization was upregulated by an average of more than 7-fold in the leaf nodule. However, lack of conservation in other strains suggests that utilization of D-galactonate may not be essential for growth of *O. dioscoreae* in leaf nodule symbiosis (Figure 4).

**Figure 4:**
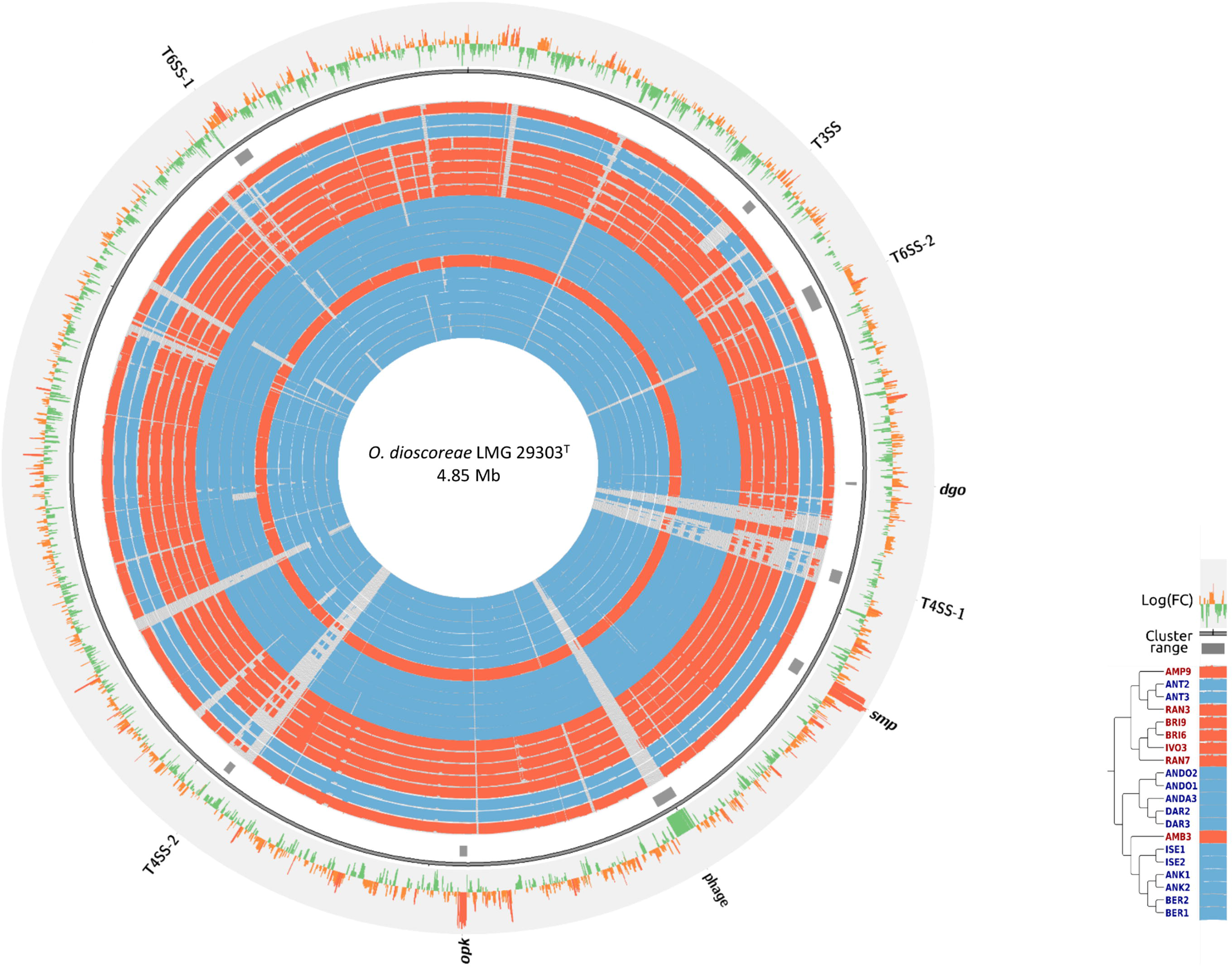
Conservation and expression of *O. dioscoreae* genes. Inner circles represent nucleotide conservation (% Blastn identity) of that genome with the type strain LMG29303^⊤^ as histograms. A legend showing the order of the genomes used for comparison is shown at the bottom right. Blue and red rings correspond to samples collected in the Diana and Atsinanana regions, respectively. Outer rings shows the log_2_ fold change of predicted genes in the leaf nodule compared to in culture as measured by RNA-Seq. Loci discussed in the text are indicated with grey rectangles.

### Simple metabolic needs of leaf nodule bacteria

Components of the GS/GOGAT pathway (ODI_R0289 – ODI_R0288; OD_R2281), were only slightly upregulated (< 3-fold) in the leaf nodule and a putative ammonium transporter (ODI_R2565) was not differentially regulated *in planta* compared to growth with ammonium as a sole nitrogen source. Neither the assimilatory nitrate reductase (ODI_R3120) or the nitrite reductase (ODI _R2365-2367) were differentially regulated. These observations indicate that nitrogen is taken up as ammonium or through deamination of amino-acids. Amino-acid biosynthetic pathways were either slightly upregulated or not differentially regulated *in planta*, with the exception of the pathways for the biosynthesis of branched-chain amino acids (Table S4) which were significantly downregulated. Several putative branched-chain amino acid transporter were simultaneously upregulated, suggesting that valine, leucine and/or isoleucine are abundant in the nodule. A *metE* homolog, coding for the cobalamin-independent methionine synthase (ODI_R2167), is upregulated by more than 150-fold on average, indicating that this pathway is preferred over the MetH pathway (ODI_R0578, not differentially regulated). LMG 29303^⊤^ cultures grew in the presence of low concentrations of yeast extract, which contains small amounts of vitamins and cofactors. The vast majority of genes (64%) involved in these pathways were not differentially regulated in the nodule. Only genes involved in vitamin B6 biosynthesis showed moderately increased expression (4.5-fold) compared to axenic cultures and may reflect a higher demand for the pyridoxal phosphate coenzyme, for example for the transamination reactions required by increased demand for amino-acid biosynthesis.

### Symbiont response to light cycle

The leaf acumen hosting the bacterial glands is composed of green tissue, raising the possibility that *O. dioscoreae* participates to or complements the metabolism of photosynthates. To test this, we generated transcriptome data of whole leaf nodules harvested in the middle of the light and dark phases. Only 8 *O. dioscoreae* genes were upregulated under light conditions (3.7-fold average), and 1 hypothetical gene was downregulated (3-fold). Six of the seven upregulated genes (ODI_R3721-3727) belonged to a putative operon conserved in all *O. dioscoreae* genomes. Aside from putative regulatory proteins and a putative ECF sigma factor, the operon encodes a short chain dehydrogenase, a flavin-containing amine oxidase, a hypothetical protein and a cyclopropane fatty acyl transferase (ODI_R3722-3725). Homologs of these, sharing on average 30-40% identity at the protein level, are induced by singlet oxygen in the purple bacterium Rhodobacter sphaeroides and may be involved in detoxifying reactive oxygen species (ROS) during photosynthesis [49]. Together, these data indicate that leaf nodule bacteria do not play a significant role in autotrophic metabolism, and reveal that detoxification of singlet oxygen, a by-product of photosynthesis, may be a significant challenge for leaf endophytes.

### The leaf nodule is dedicated to secondary metabolite production and exchange

Thirty-eight of the fifty most upregulated genes in the leaf nodule belonged to only 2 gene clusters. The largest cluster, called *smp* for its likely role in <u>s</u>econdary <u>m</u>etabolite <u>p</u>roduction comprises 23 genes upregulated by 165- to 720-fold in the nodule and is organized in 2 divergent operons, *smp1* (ODI_R1487-1493) and *smp2* (ODI_R1497-1509), separated by a regulatory region (supplementary information). Transcripts of the *smp* cluster have among the highest absolute abundance values in the leaf nodule, making up between 23.5% and 27.4% of all non-rRNA reads. The *smp* cluster is highly conserved in all *O. dioscoreae* genomes with average nucleotide identities ranging from 96.6 to 99.3% but is otherwise unique within the Alcaligenaceae. Despite this exclusivity, we could not detect any of the common signs associated with recent acquisition by horizontal gene transfer: the average %G+C of the *smp* cluster is only slightly higher than genome average (69.0% vs. 67.4%) and codon usage does not significantly deviate from the rest of the genome (Pearson chi-square test p-value > 0.25). Further, we could not detect evidence of mobile elements flanking the *smp* cluster and the phylogeny of individual *smp* genes tracked that of the whole genome (data not shown). Sequence analysis of the *smp* genes revealed a highly unusual arrangement of genes linked to non-ribosomal peptide synthesis (NRPS), putative NRPS tailoring enzymes and genes involved in acyl chain biosynthesis (Supplementary information). Iron-chelating activity on reporter medium was unchanged for an smpD null mutant, ruling out a possible role of *smp* in siderophore synthesis (supplementary information). A second gene cluster of 19 predicted genes linked to polyketide synthesis accounts for another 11 of the top 50 most differentially expressed genes. Genes of the *opk* (for *Orrella* polyketide) cluster are also among the most highly expressed in the leaf nodule, making up between 8.8% and 9.3% of all mRNA reads mapped. As for the *smp* cluster, the *opk* cluster is unique to *O. dioscoreae* among *Alcaligenaceae* but displays partial similarity to uncharacterized gene clusters of *Burkholderia glumae* BGR1 and *B. ubonensis* MSMB818. All *opk* genes appear conserved in both *Burkholderia* strains, but the homologous *Burkholderia* gene clusters contain additional genes coding for an iron containing redox enzyme and a acyl-coA dehydrogenase. Sequence analysis of the *opk* genes reveal the presence of a set of genes linked to polyketide synthesis, a class of natural products with broad activities [50], as well as an ATP transporter highly upregulated *in planta* (Supplementary information).

### Recognition of symbiotic partners and maintenance of the symbiosis

We previously reported that the genomes of *Burkholderia* leaf nodule symbionts of *Psychotria* and *Ardisia* did not encode functions found in other plant symbioses such as Type III or IV secretion, plant hormone metabolism or Nod factor synthesis [10–12]. How these symbionts avoid triggering plant defences during symbiosis remains unknown. The genome of *O. dioscoreae* LMG 29303^⊤^ encodes a type III secretion system (T3SS) (ODI_R0595-0615) and an ACC-deaminase (ODI_R1068), possibly involved in modulation of the ethylene defense pathway [51]. However, neither pathways were differentially regulated *in planta*. Furthermore, homologs of the T3SS are absent from the genomes of strains RAN3, ANT2 and ANT3 (Figure 4). Similarly, two putative type IV secretion systems (ODI_R1296-1315 and ODI_R2705-2718) are downregulated in the leaf nodule and are not conserved in all genomes. T3SS and T4SS are thus unlikely to play major roles in the interaction with the host. Two distinct type VI secretion systems (T6SS), T6SS-1 and T6SS-2 (ODI_R3980-4005, ODI_R0780-0812, respectively), were highly upregulated *in planta* and conserved in all *O. dioscoreae* genomes (Figure 4). Both clusters contain all 13 core components (TssA – TssM) and display a similar organization but with overall low sequence identity, ruling out recent duplication. Four Rhs-VgrG effectors (three in T6SS-1, one in T6SS-2) are conserved in all strains, while one effector is only conserved in ANT2 and ANT3 genomes. Additionally, an Rhs-core domain containing gene was found only in the BER1 and BER2 genomes.

### Genes under positive selection

In the absence of evidence for relaxed purifying selection affecting many vertically-transmitted symbionts, adaptive mutations may still accumulate in distinct populations of *O. dioscoreae* [52]. We found 91 genes which showed evidence of positively selected sites in *O. dioscoreae* (Table S6). Among these, 2 genes of T6SS-1 seem to be under positive selective pressure. These code for a TssA homolog (ODI_R4005) and a TssM homolog (ODI_R3983), which have respectively been proposed to be part of the T6SS baseplate and the membrane complex [53, 54]. One of four conserved VgrG effector proteins in the *O. dioscoreae* genome (ODI_R0793) also contains multiple sites with *d_N_/d_S_* > 1. Gene ODI_R3363, coding for a catalase and highly expressed in the leaf nodule, contains 3 sites under positive selection. Catalase is an important enzyme mitigating damage that may arise from ROS such as H_2_O_2_. ROS is often produced by plants in response to pathogen infection, but also plays an important role in plant signalling. Catalase has also been shown to be involved in host defence evasion by *H. pylori* by binding vitronectin [55]. Supporting this view, the sites under positive selection localize within an immune responsive domain at the C-terminal of the predicted protein, important for by T-cell interaction with *H. pylori* [56]. Together, these suggest that environmental factors such as microbe-microbe competition, but also adaptation to host immunity may be significant factors shaping the evolution of *O. dioscoreae*. Intriguingly, five genes of the *smp* cluster (ODI_R1487, ODI_R1489, ODI_R1490, ODI_R1498, ODI_R1507) display signs of positive selection, indicating that secondary metabolism may be diversifying in *O. dioscoreae* (Table S6).

## DISCUSSION

### Recent evolution of a vertically-ransmitted symbiosis in plants

Heritable symbiosis is relatively common in the animal kingdom but affects only a handful of plants [37, 57]. We describe here a new heritable symbiosis between bacteria of the *Alcaligenaceae* family and *D. sansibarensis*, the first of its kind in monocots. These symbiotic bacteria are found within conspicuous galls or nodules on leaves and are transmitted vertically via bulbils. Vertical Transmission of *O. dioscoreae* likely relies on the colonization of lateral buds, which give rise to already colonized bulbils [58]. In support of this hypothesis, high titers of *O. dioscoreae* are found in apical and lateral buds as well as bulbils and are sufficient to ensure colonization of seedlings under gnotobiotic conditions. A bacterial population maintained in proximity of developing shoot meristems, may allow both the allocation of founder colonies to the developing leaf nodules and the transmission to the next generation (Figure 5). We speculate that bacteria whose fate are either to fulfil the symbiotic function in the leaf nodule or to be transmitted to the next generation are drawn from this common pool. This unique mode of infection may have important consequences for the evolution of the symbiosis in plants. However, partial congruence between host and symbiont phylogenies suggests a mixed mode of transmission at population level, combining vertical transmission and occasional horizontal or host switching events [59]. Because even very rare horizontal transmission would completely degrade the correlation between host and symbiont phylogenetic signals [60], the vertical mode of transmission is probably largely dominant in the *D. sansibarensis*/*O. dioscoreae* symbiosis. Rare horizontal transmission may occur by host switching may occur via insect vectors, since the only evidence for a possible reservoir of environmental *O. dioscoreae* is the gut of sap feeding insects.

**Figure 5:**
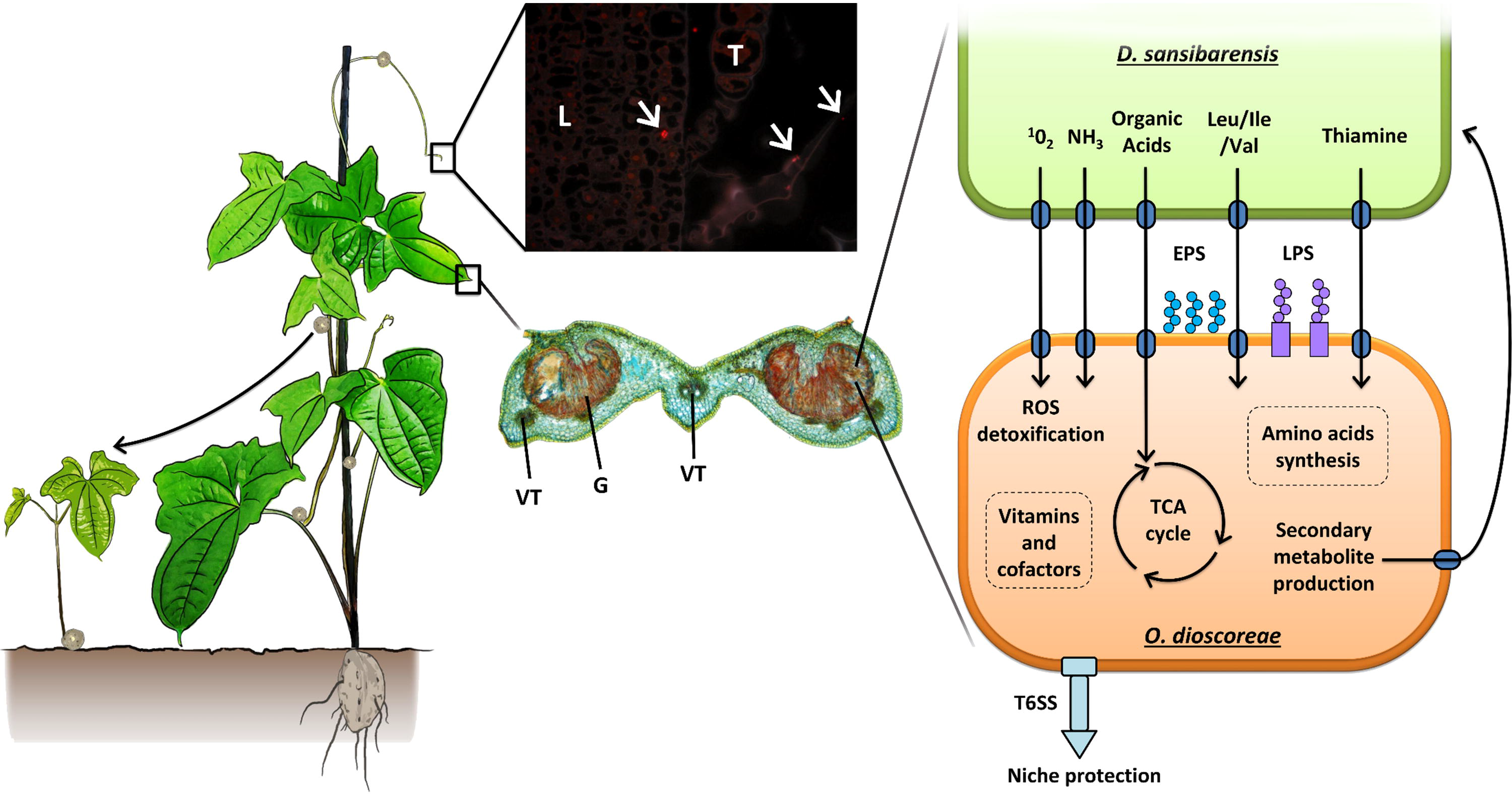
General overview of the *O. dioscoreae* lifecycle and function. *D. sansibarensis* harbors symbiotic bacteria (*O. dioscoreae*) which are contained within leaf nodules, bulbils and apical or axillary buds. The apical bud is the site of post-embryonic growth and gives rise to new leaves or aerial bulbils used for propagation. Images center: Microscopic images of *O. dioscoreae* in key plant tissues. Top center: *O. dioscoreae* in the apical bud labeled with FISH probe BETA42a specific for β-proteobacteria; L: young leaf; T: trichome; white arrows: bacteria (T. Acar, unpublished). Bottom center: Leaf nodule section triple stained with acridine red, chrysoidine and astra blue. Bacteria are often found in close proximity to trichomes, but never inside plant cells. G: bacterial gland; VT: vascular tissue. Right panel: pathways or functions presumed important for symbiosis and survival inside the leaf nodule. ROS, reactive oxygen species; LPS, lipopolysaccharide; EPS, exopolysaccharide, TCA, tricarboxylic acid cycle; T6SS, Type 6 secretion system. LPS and EPS may play a role in host/symbiont recognition. T6SS may play a role in preserving the specificity of the association by excluding competitors. See main text for details.

Vertical transmission is thought to be a particularly effective mechanism to enforce cooperation in symbiotic associations [61–63], but is not synonymous with mutualism [57, 64]. Transcriptomics analysis of leaf nodule contents show that *O. dioscoreae* relies on organic acids for growth *in planta* and scavenges iron by upregulating iron-acquisition pathways, while the pathways for biosynthesis of amino acids, as well as vitamins were expressed to levels slightly higher or comparable to growth on minimal medium. This is in direct contrast to plant pathogens, which typically show increased breadth of substrate utilization and suppression of siderophore synthesis during infection [65–67]. These data indicate that the metabolic needs of *O. dioscoreae* in the leaf nodule are relatively simple and avoid exploitation of complex host resources such as sugars, complex carbohydrates and amino-acids. Further supporting a mutualistic rather than parasitic lifestyle, T3SS genes which are essential virulence factors for a majority of gram-negative plant pathogens [68], were not significantly expressed in the leaf nodule and were not conserved in all *O. dioscoreae* genomes.

We did not find evidence for roles of the bacterial symbiont in carbon cycling, or other roles commonly associated with beneficial endophytic bacteria such as nitrogen fixation or hormone metabolism. Moderate expression of vitamin or amino-acid biosynthetic pathways, mostly consistent with growth on simple carbon and nitrogen sources, makes nutrient supply to the host unlikely. Instead, over 30% of transcripts in leaf nodule bacteria stemmed from only two gene clusters related to bacterial secondary metabolism. The role and nature of the secondary metabolites produced in the *D. sansibarensis*/*O. dioscoreae* symbiosis remains to be elucidated, but appear to result from a very unusual combination of NRPS, fatty acid and polyketide synthesis, possibly representing new molecules. We have previously shown that more than 10% of the proteome of *Ca*. *Burkholderia kirkii*, the obligate symbiont of *Psychotria kirkii*, was dedicated to the synthesis of cyclitol compounds with insecticidal an herbicidal properties [69, 70]. Similarly, the leaf nodule symbiont of *Ardisia crenata* synthesizes FR900359, a depsipeptide inhibitor of mammalian Gq proteins and potent insecticide [10, 14]. Taken together, these data suggest that leaf nodule symbionts were recruited independently in three plant families to complement the host’s secondary metabolism. Interestingly, the lack of evidence for gene conversion or horizontal transfer of the *smp* and *opk* clusters indicates that these genes were present in the last common ancestor of leaf nodule *O. dioscoreae*. Synthesis of secondary metabolites may thus be a pre-requisite for symbiont capture in the phyllosphere. Further characterization of the metabolites of the *D. sansibarensis* leaf nodule symbiosis will be essential to uncover the ecological role of this symbiosis and may provide new leads with biological activities of interest.

Vertical transmission of symbionts is a major evolutionary transition which enables the fixation of complex heritable traits in a lineage but often comes at the cost of co-dependence between the partners, with the microsymbiont unable to replicate outside of its host and in some extreme cases with the hosts unable to survive without their symbionts [57, 63, 71]. *O. dioscoreae* is among the very few vertically transmitted bacterial symbionts which can be cultured, along with symbionts of fungi [72], the Tsetse fly [73] and earthworms [74]. Unlike these examples, the genome of *O. dioscoreae* seem completely devoid of any of the hallmarks of reductive genome evolution. This unusual characteristic may be due to a very recent evolution of the association and a large effective population. The strong purifying selection acting on the *O. dioscoreae* genome would suggest the latter, but large populations would also be expected to result in high intra-host symbiont diversity [34, 63, 75, 76]. Surprisingly, we found intra-host diversity to be very low and consistent with accumulation of low frequency *de novo* mutations. Perhaps as another consequence of the unique infection mode directly coupled to the host’s post-embryonic growth, intra-host competition between genotypes may instead account for this low observed diversity [77]. Because host controls such as sanctions have never been documented in heritable symbioses [62, 78, 79], hypothetical mechanisms preventing the fixation of non-cooperating genotypes as a result of intense intra-host competition remains to be elucidated.

How leaf nodule symbionts avoid triggering innate plant immune defenses remains an open question. We show that T3SS and T4SS are not conserved in *O. dioscoreae* genomes, suggesting that these secretion systems do not play an active role in modulating the immune response of the host. The genomes of *O. dioscoreae* do not encode cell-wall degrading enzymes and the metabolism of organic acids rather than complex sugars may avoid the release of damage associated molecular patterns (DAMPs) [80]. Alternatively, EPS, upregulated in the leaf nodule, is known to be crucial for the establishment of successful symbiosis with legumes [81, 82] and may play an important role in protection against host defence [83]. Interestingly, we could also detect signatures of positive selection in natural populations of *O. dioscoreae* in various genes coding for products implicated in the elicitation of plant defenses including flagella, several GGDEF domain proteins and proteins involved in iron uptake and siderophore synthesis [84]. These potential M/PAMPs may have evolved to lower immune recognition and enable the bacteria to multiply within host tissue (e.g. in the shoot apical bud) without triggering defenses. Specific epitopes may also enable active recognition of beneficial symbionts, similar to what has been described in some invertebrate systems such as *Hydra* sp. or in the squid-*Vibrio* symbiosis [85, 86]. In this respect, patterns of positive selection on *smp* genes are of particular interest. Diversifying selection on genes of the secondary metabolism may reflect adaptation to biotic pressure in distinct environments, e.g. the presence of different herbivores, but may also reflect pressure by the host to indirectly gauge the contribution of the leaf nodule symbionts. These genes make attractive targets for mutagenesis or heterologous expression to further elucidate partner recognition in leaf nodule symbiosis.

Because the lifecycle of *O. dioscoreae* is strictly extracellular and leaf nodules initially appear open to the outside environment [16], it is unclear how specificity is maintained in the leaf nodule. A complex interplay of immune modulation or recognition and filtering due to environmental parameters may enable the host to control access to the nodule. Oxidative stress, possibly as a result of singlet oxygen produced through photosynthesis, may guard against potential invading microorganisms [87]. In addition, we found two conserved *O. dioscoreae* T6SS to be upregulated in the leaf nodule. T6SS is a common mediator of antagonistic microbe-microbe interactions in many bacteria and may actively contribute to gatekeeping inside the leaf nodule [88]. Evidence of positive selection in genes of the T6SS, including diversification of putative effectors, could indicate ongoing adaptation of the symbiont to maintain its niche against biotic challenges. This phenomenon was recently observed in bee gut symbionts and was hypothesized to have a significant impact into shaping the evolution of the bee gut microbiome [89].

In conclusion, this study broadens our knowledge of plant-bacteria interactions in the phyllosphere and highlights features specific to heritable symbioses in plants. We demonstrate striking commonalities between the leaf nodule symbiosis in *D. sansibarensis* and those of Rubiaceae and Primulaceae, including a vertical mode of symbiont transmission and a seemingly central role of bacterial secondary metabolism. Because *O. dioscoreae* can easily be cultured, and the host plant can easily be grown and propagated using standard methods, this binary symbiosis provides an attractive model system for the study of beneficial plant-bacteria interactions in the phyllosphere.

## ACKNOWLEDGMENTS

The authors wish to thank Dr. Olivier Leroux (Ghent University) for his help with microscopy. This work was supported by the Flemish Fonds Wetenschappelijk Onderzoek under grant G017717N and the Special Research fund of Ghent University under grant BOF17/STA/024. The funders had no role in study design, data collection and analysis, decision to publish, or preparation of the manuscript.

