## supplementary figures and tables for "Early steps in the evolution of vertical transmission revealed by a plant-bacterium symbiosis"

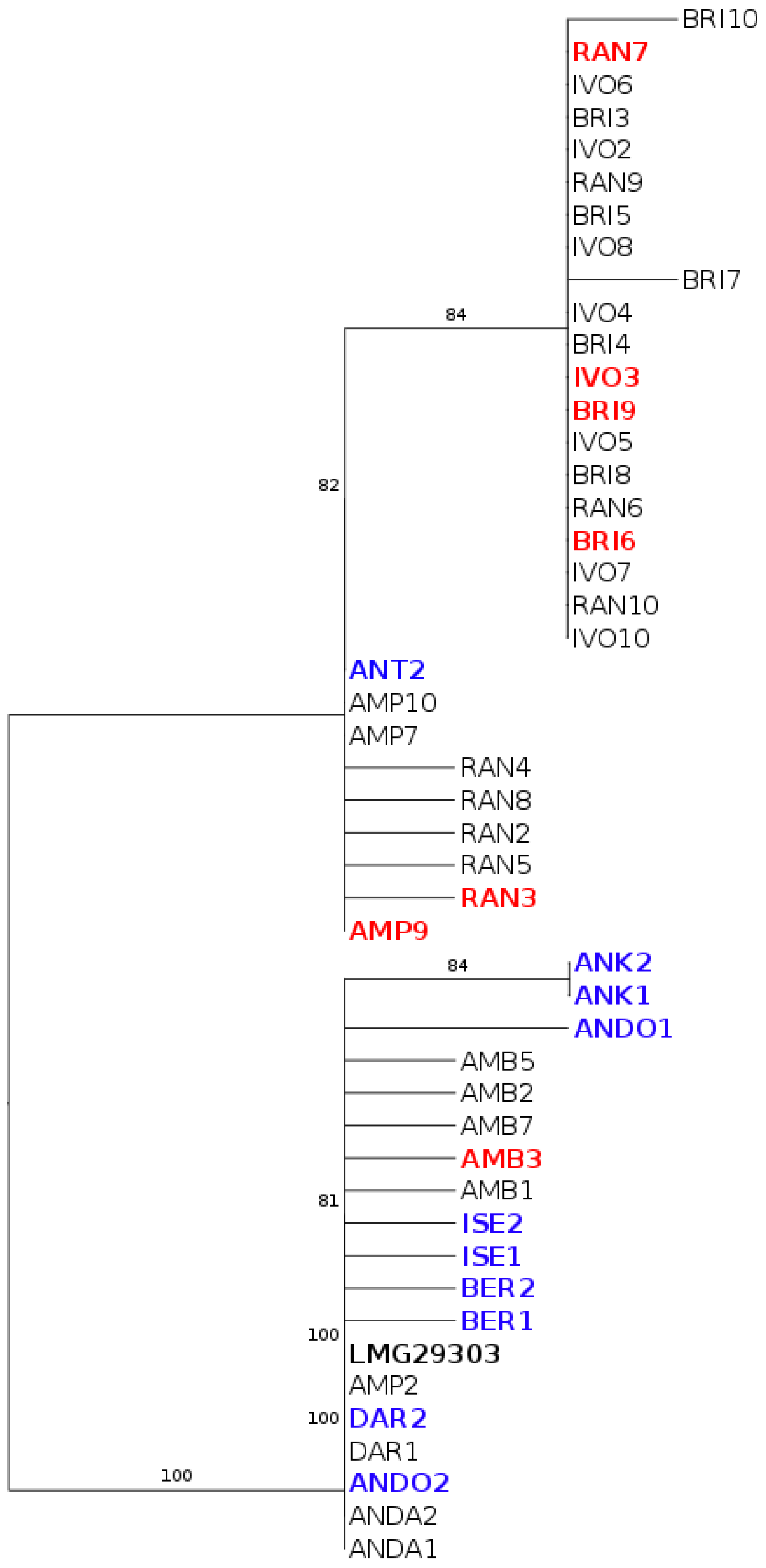


**Figure S1**: NrdA gene phylogeny with bootstrap values from wild-collected *O. dioscoreae.* Colored sample names represent samples sent for WGS. Red: samples from the east region; Blue: samples from the north region

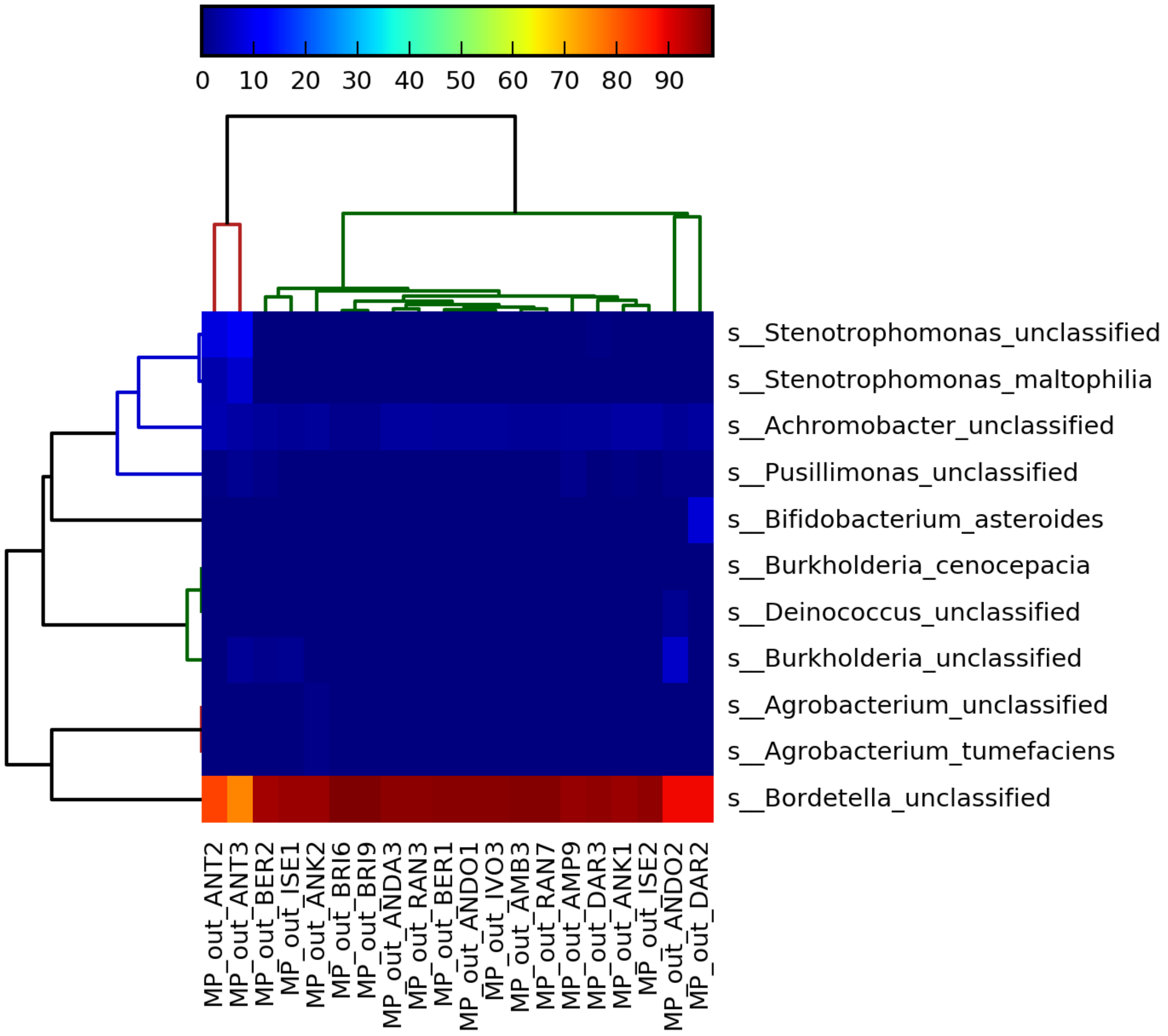


**Figure S2**: Heatmap of OTU abundance in collected *D. sansibarensis* leaf nodules as computed by MetaPhlan

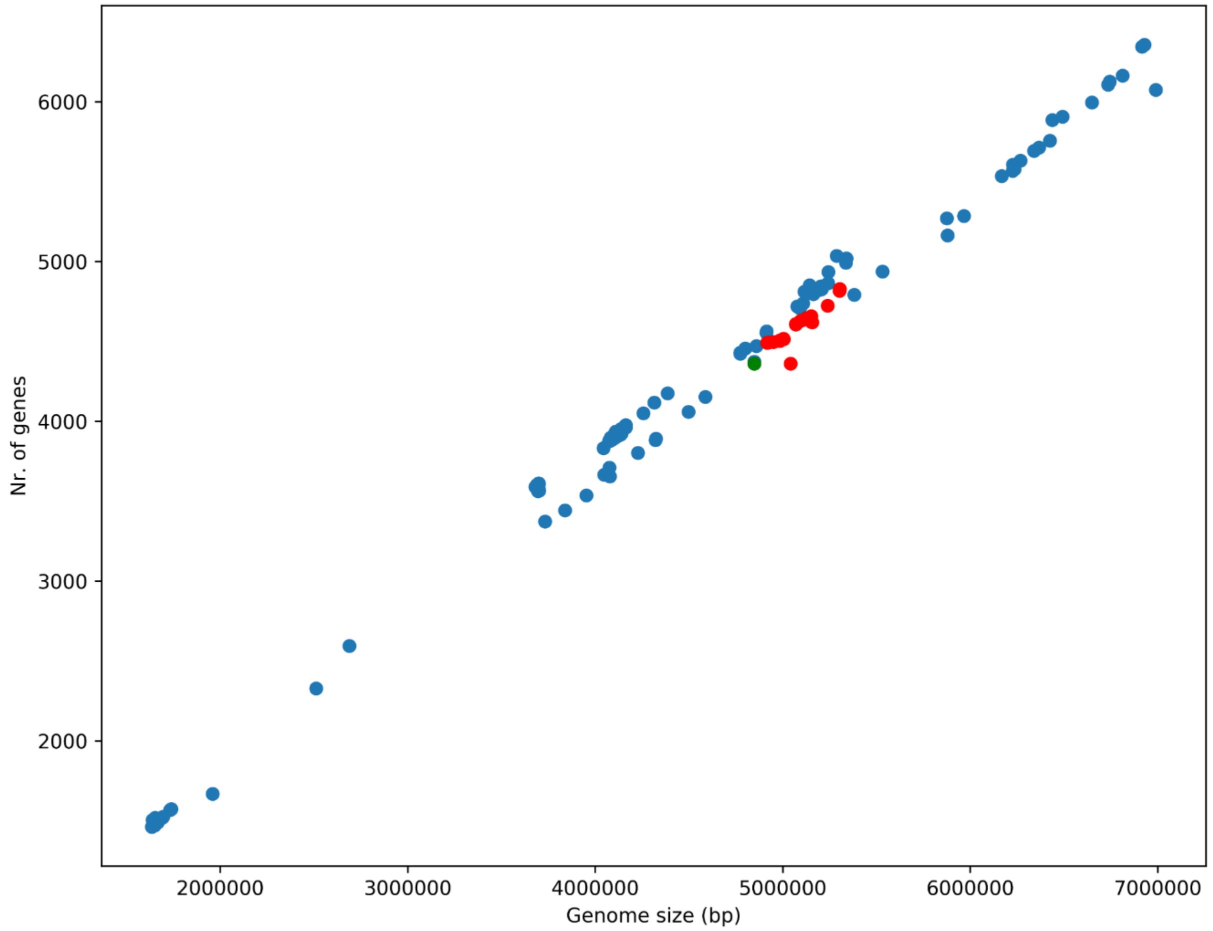


**Figure S3**: Genome size and number of genes of bacteria in the Alcaligenaceae family. RefSeq assemblies of all species belonging to the Alcaligenaceae family (taxonomy id 506, n=449) were downloaded from Genbank (accessed 13-11-2018) Green: *O. diosocoreae* typestrain (LMG29303$^{T}$) Red: *O. dioscoreae* strains from Madagascar.

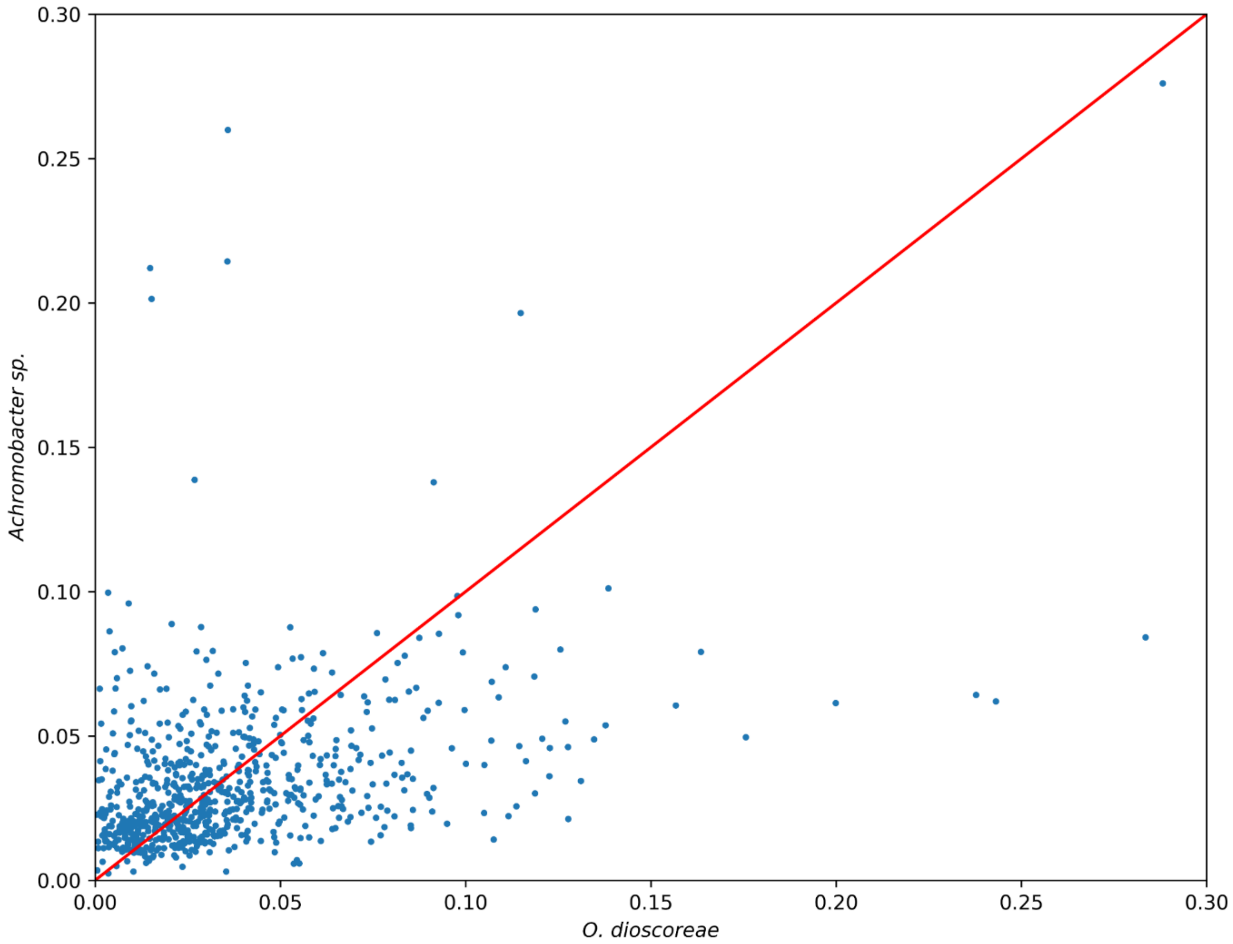


**Figure S4**: Comparison of the ratio of synonymous (dN) and nonsynonymous (dS) substitutions of the single-copy orthologs of several *Achromobacter sp.* (ENA accessions: ASM23678v2, ASM21974v1, GCS2v1, ASM118959v1, ASM163968v1, ASM16583v1, Achr_xylo_C54_V2, ASM105105v1) and sampled *O. dioscoreae* genomes. Zero values were removed from the analysis. Mean *O. dioscoreae* dN/dS (0.0374) and mean *Achromobacter sp.* dN/dS (0.0339) did not differ significantly (Wilcoxon rank-sum p-value: 0.85)

**Table S1: Strains and plasmids used in this study**

| Strain | Genotype/Description | Source/Reference |
|---|---|---|
| ***O. dioscoreae* strains** | | |
| LMG 29303T | Leaf nodule isolate - Switzerland | (Carlier et al., 2017) |
| R67088 | Leaf nodule isolate - Belgium | (Carlier et al., 2017) |
| R67584 | Leaf nodule isolate - Congo | This Study |
| LF01 | LMG 29303T, $Nal^R$ | This Study |
| FID1 | LF01 △smpD, $Km^R$ | This Study |
| ***E. coli* strains** | | |
| Top 10 | | Invitrogen |
| S17-1 | Donor, recA pro hsdR RP4-2-Tc::Mu-Km::Tn7 integrated into the chromosome | (Simon R et al., 1983) |
| **Plasmids** | | |
| pDONRpEX18 | Gateway adapted donor vector; attP1 and attP2, sacB, $Te^R$ | (Hmelo et al., 2000) |
| pKD4 | $Km^R$-cassette | (Datsenko & Wanner, 2015) |

**Table S2.** ***Dioscorea sansibarensis*** **sampling locations in Madagascar.**

| Identifier | Locality | GPS coordinates | | Date of sampling |
|---|---|---|---|---|
| ANDA | Andranomanitra | S12°25’10.3’’ | E49°20’40.4’’ | 05/2017 |
| DAR | Daraina | S12°29’07.1’’ | E49°23’46.4’’ | 05/2017 |
| ANT | Antsakoabe | S12°39’48.1’’ | E49°17’42.7’’ | 05/2017 |
| ANDO | Andranonakoko | S12°55’36.8’’ | E49°11’27.7’’ | 05/2017 |
| ISE | Isesy | S13°08’44.8’’ | E49°04’49.1’’ | 05/2017 |
| BER | Beramanja | S13°22’14.0’’ | E48°52’18.1’’ | 05/2017 |
| ANK | Ankazomahity | S13°28’56.8’’ | E48°43’42.4’’ | 05/2017 |
| BRI | Brickaville | S18°49’19.1’’ | E49°04’36.5’’ | 11/2016 |
| AMB | Ambavaniasy | S18°56’42.9’’ | E48°30’48.0’’ | 11/2016 |
| RAN | Ranomafana | S18°58’35.7’’ | E48°55’04.8’’ | 11/2016 |
| IVO | Ivoala | S19°02’21.3’’ | E48°55’59.6’’ | 11/2016 |
| AMP | Ampaho | S19°26’56.1’’ | E48°55’18.2’’ | 11/2016 |

**Table S3. Oligonucleotide primers used in this study**

| Primer name | Sequence (5' - 3') |
|---|---|
| **Primers used for phylogeny** | |
| Odio_nrdA_R | GAGTCCTTCGCCTTCTTG |
| Odio_nrdA_F | ACTACTTCCCCACCTTCA |
| matK-sans-R | CTAGCACACGAAATCCGAA |
| matK-sans-F | AATACCCCATCCCATCCA |
| rbcL-sans-R | CGCGATGGATGTGAAGAA |
| rbcL-sans-F | TACGTGGTGGACTTGATTTT |
| rpl32-trnL-R | ATTTGGAAGAAAAGGGGGT |
| rpl32-trnL-F | TTCCTAAGAGCAGCGTGT |
| **Primers used for quantitative PCR** | |
| rpoD-06-R | CACCATTTCGGTCAGCTT |
| rpoD-05-F | GAACTCCCCCGCAAACAA |
| galactonate_dehydratase-02-R | CTCCTTCACGTACTCCTC |
| galactonate_dehydratase-01-F | CTACAACGCCTTCATCCA |
| KASII-02-R | AAGAACAGGCCATCGACA |
| KASII-01-F | GAGATCGCGGAAAACCAG |
| PqqC-02-R | TGAAGAACACCACCGCCT |
| PqqC-01-F | CTACCGCGACATGCTCAA |
| ImpA-01-R | TCACGCAGGGACAATTGG |
| ImpA-02-F | GCAGCAGTATTGGGAAGG |
| nrp-02-R | GCGAAGTTGAAGGTATAGGG |
| nrp-01-F | GGTGTTCGCTGCCTATTG |
| **Primers used for smpD-mutant construction** | |
| 00690-Dn-GW-03-F | TACAAGAAAGCTGGGTAGACGGAATCCAGCCAGAC |
| 00690-Dn-kan-04-R | CGGAATAGGAACTAAGGAGGATATTCATATGGCAAGTCATGCAACACCAGA |
| 00690-Up-kan-05-R | GAACTTCGAAGCAGCTCCAGCCTATCGTGATAGACATGGGAGGA |
| 00690-Up-GW-06-F | TACAAAAAAGCAGGCTTCCGTGTTGCAAGAGGAGAT |
| pKD4rev-2 | CATATGAATATCCTCCTTAGTTCCTATTCCG |
| pKD4fwd-2 | TAGGCTGGAGCTGCTTCGAAGTTC |

**Table S5. Biolog PM1/PM2A read-out. Purple color development was scored at 24h and 48h.**

| Substrate C-source | Growth after | |
|---|---|---|
| | **24h** | **48h** |
| Succinic Acid | x | |
| L-Proline | x | |
| L-Lactic Acid | x | |
| L-Glutamic Acid | | x |
| D-Galactonic Acid-γ-Lactone | | x |
| D,L-Malic Acid | x | |
| α-Keto-Glutaric Acid | x | |
| m-Tartaric Acid | x | |
| α-Hydroxy Glutaric Acid-γ-Lactone | x | |
| Citric Acid | x | |
| Fumaric Acid | x | |
| Propionic Acid | | x |
| Mono Methyl Succinate | | x |
| Methyl Pyruvate | x | |
| D-Malic Acid | x | |
| L-Malic Acid | x | |
| Pyruvic Acid | x | |
| Citraconic Acid | | x |
| β-Hydroxy Butyric Acid | x | |
| Oxalomalic Acid | x | |
| Succinamic Acid | x | |
| D-Tartaric Acid | x | |
| L-Pyroglutamic Acid | x | |

**Table S6. List of positive selected genes**

| Gene | Nr. of sites under positive selection | Average dN/dS | Log2FC | Percentile expression | Gene product | Localization |
|---|---|---|---|---|---|---|
| ODI_R0217 | 1 | 8.32 | -1.33 | 24.31 | Predicted transcriptional regulator LiuR of leucine degradation pathway, MerR family | Cytoplasmic |
| ODI_R0262 | 1 | 5.12 | 0 | 29.54 | Outer membrane stress sensor protease DegS | Periplasmic |
| ODI_R0275 | 1 | 6.87 | -3.17 | 24.81 | Histidinol dehydrogenase | Cytoplasmic |
| ODI_R0308 | 1 | 6.89 | 2.16 | 5.41 | Cytochrome c oxidase polypeptide I | CytoplasmicMembrane |
| ODI_R0387 | 1 | 6.54 | 0 | 12.63 | hypothetical protein | Unknown |
| ODI_R0475 | 1 | 4.13 | -0.93 | 37.82 | hypothetical protein | Unknown |
| ODI_R0485 | 3 | 7.23 | -0.54 | 23.92 | Hypothetical oxidoreductase, YbiC homolog | Cytoplasmic |
| ODI_R0541 | 3 | 9.39 | -2.11 | 7.65 | hypothetical protein | Unknown |
| ODI_R0556 | 2 | 7.45 | -1.89 | 47.67 | Phosphate regulon sensor protein PhoR (SphS) | CytoplasmicMembrane |
| ODI_R0578 | 1 | 2.84 | 1.23 | 10.5 | 5-methyltetrahydrofolate--homocysteine methyltransferase | Cytoplasmic |
| ODI_R0643 | 1 | 4.33 | -1.9 | 24.17 | Xanthine and CO dehydrogenases maturation factor, XdhC/CoxF family | Cytoplasmic |
| ODI_R0690 | 1 | 4.74 | 0.69 | 45.63 | Probable transmembrane protein | CytoplasmicMembrane |
| ODI_R0698 | 6 | 8.99 | 2 | 36.4 | hypothetical protein | Unknown |
| ODI_R0813 | 1 | 6.86 | -1.22 | 20.87 | hypothetical protein | Unknown |
| ODI_R0862 | 3 | 7.31 | 1.54 | 24.72 | Short chain dehydrogenase | Extracellular |
| ODI_R0900 | 1 | 6.8 | 0.52 | 1.63 | 18K peptidoglycan-associated outer membrane lipoprotein; Peptidoglycan-associated lipoprotein precursor; Outer membrane protein P6; OmpA/MotB precursor | OuterMembrane |
| ODI_R0918 | 2 | 5.03 | 0 | 22.34 | Tricarboxylate transport protein TctB | CytoplasmicMembrane |
| ODI_R0946 | 1 | 5.74 | 0 | 43.89 | D-alanine--D-alanine ligase | Cytoplasmic |
| ODI_R0990 | 2 | 4.84 | 1.04 | 13.92 | GGDEF domain protein | Cytoplasmic |
| ODI_R1014 | 1 | 5.02 | 0.95 | 17.02 | TRAP-type C4-dicarboxylate transport system, large permease component | CytoplasmicMembrane |
| ODI_R1071 | 4 | 8.76 | -0.33 | 34.68 | Acetylornithine deacetylase/Succinyl-diaminopimelate desuccinylase and related deacylases | Cytoplasmic |
| ODI_R1118 | 2 | 8.46 | 2.51 | 28.13 | 4-hydroxy-tetrahydrodipicolinate reductase | Cytoplasmic |
| ODI_R1119 | 1 | 6.91 | 3.44 | 18.4 | Outer membrane lipoprotein SmpA, a component of the essential YaeT outer-membrane protein assembly complex | OuterMembrane |
| ODI_R1148 | 2 | 8.65 | 1.94 | 28.2 | FIG152265: Sodium:solute symporter associated protein | CytoplasmicMembrane |
| ODI_R1156 | 1 | 4.79 | -1.11 | 38.21 | Methylglutaconyl-CoA hydratase | Cytoplasmic |
| ODI_R1187 | 1 | 5.7 | 2.01 | 25.15 | UPF0246 protein YaaA | Cytoplasmic |

| | | | | | | |
|---|---|---|---|---|---|---|
| ODI_R1403 | 1 | 6.91 | 1.48 | 12.52 | RidA/YER057c/UK114 superfamily, group 2, YoaB-like protein | Cytoplasmic |
| ODI_R1418 | 1 | 5.84 | 2.32 | 49.34 | Tetrapyrrole methylase family protein | Unknown |
| ODI_R1419 | 1 | 7.18 | 0 | 30.56 | FIG146278: Maf/YceF/YhdE family protein | Cytoplasmic |
| ODI_R1457 | 1 | 5.91 | -1.4 | 40.02 | putative 4-hydroxybenzoyl-CoA thioesterase | CytoplasmicMembrane |
| ODI_R1487 | 1 | 5.85 | 7.64 | 1.11 | Thioesterase in siderophore biosynthesis gene cluster | Cytoplasmic |
| ODI_R1489 | 1 | 3.02 | 8.11 | 0.27 | FIG00663483: hypothetical protein | Cytoplasmic |
| ODI_R1490 | 2 | 3.95 | 8.14 | 0.07 | Siderophore biosynthesis non-ribosomal peptide synthetase modules | CytoplasmicMembrane |
| ODI_R1498 | 1 | 8.21 | 8.16 | 0.16 | Tricarboxylate transport protein TctC | CytoplasmicMembrane |
| ODI_R1507 | 5 | 7.29 | 8.51 | 0.72 | 4'-phosphopantetheinyl transferase | Unknown |
| ODI_R1513 | 1 | 6.86 | 0.86 | 23.4 | transcriptional regulator RhlR | Cytoplasmic |
| ODI_R1522 | 1 | 7.61 | 3.47 | 6.9 | Cytidylate kinase | Cytoplasmic |
| ODI_R1692 | 2 | 6.63 | 0 | 33.18 | tRNA-specific 2-thiouridylase MnmA | Cytoplasmic |
| ODI_R1761 | 2 | 6.71 | 2.35 | 44.7 | Ferric siderophore transport system, biopolymer transport protein ExbB | CytoplasmicMembrane |
| ODI_R1795 | 1 | 5.54 | 1.48 | 27.05 | Phosphopantothenoylcysteine decarboxylase / Phosphopantothenoylcysteine synthetase | Cytoplasmic |
| ODI_R1877 | 1 | 8.63 | -0.54 | 27.46 | Ribosome-associated heat shock protein implicated in the recycling of the 50S subunit (S4 paralog) | Cytoplasmic |
| ODI_R1904 | 1 | 6.36 | 1.74 | 22.41 | FIG053235: Diacylglucosamine hydrolase like | Cytoplasmic |
| ODI_R1905 | 2 | 5.67 | 2.06 | 23.43 | Glutamate-ammonia-ligase adenylyltransferase | Cytoplasmic |
| ODI_R1920 | 1 | 8.62 | -2.3 | 45.09 | Transcriptional regulators | Unknown |
| ODI_R1924 | 1 | 7.57 | -2.7 | 39.7 | L-ectoine synthase | Cytoplasmic |
| ODI_R2027 | 2 | 5.96 | -2.03 | 33.16 | Transcriptional regulator, LysR family | Cytoplasmic |
| ODI_R2071 | 1 | 5.14 | -0.86 | 49.66 | Manganese ABC transporter, inner membrane permease protein SitD | CytoplasmicMembrane |
| ODI_R2086 | 1 | 6.08 | 0 | 34.93 | hypothetical protein | Unknown |
| ODI_R2106 | 1 | 7.27 | -2.05 | 23.61 | ABC transporter ATP-binding protein | CytoplasmicMembrane |
| ODI_R2287 | 2 | 6.96 | 2.38 | 18.74 | Ferric iron ABC transporter, iron-binding protein | Periplasmic |
| ODI_R2311 | 1 | 4.12 | 1.53 | 24.06 | hypothetical protein | Extracellular |
| ODI_R2324 | 1 | 5.85 | -0.52 | 43.68 | Anthranilate phosphoribosyltransferase like | Cytoplasmic |
| ODI_R2348 | 2 | 4.6 | -1.23 | 49.3 | | Cytoplasmic |
| ODI_R2356 | 1 | 6.21 | 2.79 | 39.25 | FIG139991: Putative thiamine pyrophosphate-requiring enzyme | CytoplasmicMembrane |
| ODI_R2368 | 1 | 6.47 | 0 | 49.28 | Arginine/ornithine antiporter ArcD | CytoplasmicMembrane |
| ODI_R2386 | 3 | 9.85 | -0.76 | 12.97 | Glycerol-3-phosphate ABC transporter, periplasmic glycerol-3-phosphate-binding protein (TC 3.A.1.1.3) | Periplasmic |
| ODI_R2432 | 1 | 8.47 | -0.82 | 11.27 | putative membrane protein | CytoplasmicMembrane |

| | | | | | | |
|---|---|---|---|---|---|---|
| ODI_R2541 | 1 | 6.5 | -1.15 | 46.76 | Transcriptional regulator, ArsR family | Unknown |
| ODI_R2585 | 1 | 4.21 | 2.73 | 17.63 | hypothetical protein | Unknown |
| ODI_R2603 | 1 | 7.14 | -1.13 | 19.44 | N-acetylmannosaminyltransferase | Cytoplasmic |
| ODI_R2647 | 2 | 3.9 | -1.69 | 11.5 | putative sigma-54-dependent transcriptional regulator | Cytoplasmic |
| ODI_R2650 | 1 | 6.87 | 0 | 32.82 | Permease of the drug/metabolite transporter (DMT) superfamily | CytoplasmicMembrane |
| ODI_R2651 | 2 | 4.35 | -0.5 | 6.7 | diguanylate cyclase/phosphodiesterase (GGDEF & EAL domains) with PAS/PAC sensor(s) | CytoplasmicMembrane |
| ODI_R2844 | 1 | 3.37 | 0 | 16.12 | Cyanophycin synthase | Cytoplasmic |
| ODI_R2907 | 1 | 4.9 | -2.23 | 14.46 | putative lipoprotein | Unknown |
| ODI_R2912 | 1 | 4.43 | 1.32 | 15.57 | Type II/IV secretion system protein TadC, associated with Flp pilus assembly | CytoplasmicMembrane |
| ODI_R2954 | 1 | 5.49 | 1.49 | 41.38 | Permease of the drug/metabolite transporter (DMT) superfamily | CytoplasmicMembrane |
| ODI_R3070 | 1 | 6.92 | -1.05 | 12.27 | NADPH:quinone oxidoreductase | Cytoplasmic |
| ODI_R3085 | 2 | 7.52 | -1.44 | 29.45 | FIG004556: membrane metalloprotease | CytoplasmicMembrane |
| ODI_R3122 | 1 | 5.69 | -2.64 | 42.01 | hypothetical protein | Unknown |
| ODI_R3124 | 2 | 8.02 | 2.98 | 45.77 | putative cytoplasmic protein | Unknown |
| ODI_R3158 | 1 | 6.52 | 1.33 | 42.17 | hypothetical protein | Unknown |
| ODI_R3173 | 2 | 6.42 | 0.61 | 45.36 | Survival protein SurA precursor (Peptidyl-prolyl cis-trans isomerase SurA) | CytoplasmicMembrane |
| ODI_R3287 | 1 | 7.38 | -0.38 | 39.79 | Oxygen-insensitive NADPH nitroreductase | Unknown |
| ODI_R3296 | 1 | 4.76 | 2.54 | 16.57 | Mannosyltransferase OCH1 and related enzymes | Unknown |
| ODI_R3358 | 1 | 3.91 | 0 | 48.55 | Sensor protein PhoQ | Unknown |
| ODI_R3363 | 3 | 5.72 | 4.66 | 1.77 | Catalase | Periplasmic |
| ODI_R3392 | 2 | 7.48 | 0.56 | 41.49 | Molybdenum cofactor biosynthesis protein MoaC | Cytoplasmic |
| ODI_R3422 | 2 | 6.28 | 0 | 38.93 | GCN5-related N-acetyltransferase | Unknown |
| ODI_R3551 | 1 | 5.33 | 1.94 | 14.58 | ATP-dependent DNA ligase LigC | Cytoplasmic |
| ODI_R3585 | 2 | 5.92 | -0.67 | 48.14 | NADH-ubiquinone oxidoreductase chain F | Cytoplasmic |
| ODI_R3591 | 1 | 4.58 | 1.33 | 15.05 | Proline dehydrogenase | Unknown |
| ODI_R3683 | 4 | 7.42 | -2.19 | 42.21 | FIG00858788: hypothetical protein | Cytoplasmic |
| ODI_R3983 | 1 | 4.28 | 1.52 | 32.71 | IcmF-related protein | CytoplasmicMembrane |
| ODI_R4005 | 4 | 6.07 | 4.4 | 15.21 | Uncharacterized protein ImpA | Cytoplasmic |
| ODI_R4009 | 1 | 6.41 | 3.24 | 36.1 | Methylated-DNA--protein-cysteine methyltransferase | Cytoplasmic |
| ODI_R4021 | 3 | 6.97 | 1.18 | 32.25 | Leader peptidase (Prepilin peptidase) / N-methyltransferase | CytoplasmicMembrane |
| ODI_R4056 | 1 | 4.49 | 0.67 | 9.14 | Oligopeptide transport system permease protein OppB (TC 3.A.1.5.1) | CytoplasmicMembrane |

| | | | | | | |
|---|---|---|---|---|---|---|
| ODI_R4236 | 2 | 9.21 | 0 | 44.5 | Histone acetyltransferase HPA2 and related acetyltransferases | Cytoplasmic |
| ODI_R4269 | 1 | 5.83 | 0 | 39.9 | 5-formyltetrahydrofolate cyclo-ligase | Unknown |
| ODI_R4382 | 2 | 8.8 | 0 | 28.5 | A/G-specific adenine glycosylase | CytoplasmicMembrane |
