## supplementary information for "Early steps in the evolution of vertical transmission revealed by a plant-bacterium symbiosis"

1. **DNA extraction and PCR**

Silica-dried samples were processed using a combination of bead beating (Retsch MM400, Haan, Germany) and the Maxwell® 16 DNA Purification Kit (Promega, Madison, WI, USA). Briefly, a single dehydrated sample, 567 µL TE buffer and one 6 mm glass bead (280 mg) were added to a 2 ml microcentrifuge tube and processed twice in a tissue lyser for 1 min 15 s at a frequency of 30 Hz. Homogenized samples were stored on ice for 2 minutes between each run to prevent overheating. A lysis solution (30 µL of 10% SDS, 3 μL of 20 mg/mL proteinase K) was added and incubated at 37°C for 1h. Following incubation, samples were centrifuged briefly at 14 000 rpm, and the supernatant was transferred into Maxwell® 16 DNA Purification Kit cartridges. The remainder of the DNA extraction protocol was carried out in the Maxwell® 16 Instrument according to the manufacturer’s instructions (Promega). Afterwards, RNA was removed from the gDNA with 1 μg/mL RNase A for 1h at room temperature. Species-specific primers for *nrdA* were used to confirm the presence of *Orrella dioscoreae*. For plant species identification, PCR analysis was done using primers specific for the chloroplastic markers *matK*, *rbcL* and *rpl32*-*trnL*. Sequences were aligned using Muscle [1] and manually trimmed using CLC Main Workbench v7.7.2. (Qiagen) and compared to reference sequences obtained from a vouchered *D. sansibarensis* specimen from the live collection of the botanical garden of Ghent University (accession 19001189). All specific oligos used in this study are listed in Table S3.

2. **Reference genome assembly and annotation**

The previously reported draft genome of *O. dioscoreae* strain LMG 29303^T^ was closed using PacBio long read sequencing. Genomic DNA was isolated as described previously [2] and long read libraries were constructed and sequenced using a Pacific Biosciences RSII instrument and P6/C4 chemistry on one SMRT cell (Pacific Biosciences). The sequencing run yielded 47 969 reads, with an N50 of 26 386 bp and an average read length of 19 782 bp. The reads were assembled *de novo* using the HGAPv2 program of the PacBio SMRT portal software suite with default settings, yielding 9 contigs for a total polished assembly size of 4 839 294 bp and an N50 of 2.69 Mb. Previous PFGE analysis indicated that the genome of LMG 29303T consists of a single chromosome without plasmids [2]. Contigs were further assembled into scaffolds using the SSPACE-LongRead v1-1 software [3], yielding a single scaffold with several sequence gaps. Gaps were further closed by combining scaffolded contigs, contigs obtained previously using Illumina MiSeq technology, and 2x250 bp paired-end Illumina MiSeq reads (total estimated coverage of 155.5x), using GMcloser [4]. Overlapping scaffold ends were trimmed in CLC Main Workbench v7.7.2 and the final chromosome assembly was circularized, yielding a final assembly of 4 848 101 bp. Finally, the chromosome sequence was polished using Pilon [5] and the 2x250 bp Illumina MiSeq reads used to generate the draft genome assembly. Genome sequence annotations were automatically transferred from the draft genome using the RATT tool, part of the PAGIT toolkit [6]. Annotation curation was done in Artemis [7]. The completed genome sequence of *O. dioscoreae* LMG 29303^T^ is deposited in the European Nucleotide Archive under accession number GCA_900089455.2.

3. **Chloroplast genome assembly**

Fresh leaf nodules of a specimen of *D. sansibarensis* (from which strain R-67088 was isolated) were ground with a mortar and pestle and DNA was extracted and sequenced as described above. A total of 4.7M Illumina 2x150 bp read pairs were assembled with SPAdes 3.0 in metagenome mode as above. Contigs were binned according to %G+C, average coverage and taxonomic assignment. Sequencing reads corresponding to the binned contigs were extracted and reassembled using SPAdes 3.0 in single genome mode with –k 21, 33, 55, 77, 99, resulting in a total assembly length of 459 173 bp distributed over 68 contigs and an average coverage of 24.7x. Individual contigs were searched against the NCBI RefSeq database (accessed Feb. 2017) using the Blastn program [8] with an e-value cut-off of 0.001. Only contigs with hits against plastid genome sequences were kept, resulting in a draft assembly of the *D. sansibarensis* chloroplast of 128 321 bp in 6 contigs. Average sequence coverage was 25.3x, except for a 33.5 kb contig displaying an average coverage of 55.9x corresponding to unresolved copies of the large inverted repeat common in chloroplast genomes [9]. The *Dioscorea sansibarensis* chloroplast sequence was deposited in the European Nucleotide Archive under project accession PRJEB30106.

4. **Metagenome project mining**

To investigate the presence of *O. dioscoreae* in the environment, metagenomics data from the MG-RAST [10] database was screened. Project data of 71 metagenome projects (totalling 1677 metagenome samples) was acquired using the MG-RAST toolkit, and the rRNA sequences were extracted (totalling over 85 Gb of rRNA data). Blast [8] was used to search for sequencing matching the 16S rRNA sequence of *O. dioscoreae.* Sequence hits with ≥ 98% identities were further analysed using SILVA [11], and compared to the NCBI nr dna database to determine the most likely origin of the sequence. No sequence from any of these 1677 samples from environments as diverse as soil, plants, water bodies, feces, and anthropogenic environments could be confidently assigned to the genus *Orrella* above the 98% rRNA identity threshold.

5. **Detection of horizontal gene transfer**

Presence of horizontal gene transfer was investigated using two methods. In the first method, CDS sequences of all single-copy core genes were aligned using Muscle [1], trimmed using Trimal [12], and a phylogenetic tree was created using FastTree [13]. The resulting trees were rooted using *Achromobacter xyloxidans* as an outgroup, or by midpoint rooting if no ortholog was present in *A. xyloxidans*. Trees resulting from alignments with only few divergent sequences (< 11) were discarded. The presence of incongruencies between gene and species trees was tested using the ETE3 package in Python [14]. Incongruent trees and alignment were further manually checked for possible confounding factors. The second method involved the detection of gene conversion by PhiPack [15]. P-values resulting from the phi-test for gene conversion were corrected using the Bonferroni correction for multiple testing.

6. **Measurement of free oxygen concentration**

Fresh mature leaves of *D. sansibarensis* were collected in the greenhouse immediately before the experiment. Oxygen concentration was measured 1 mm below the abaxial surface of one of the leaf gland channels closest to the mid-rib using an O_2_ microsensor (100 m tip size, Unisense). The sensor was connected to a OXY Meter microsensor amplifier (Unisense) and the data collected and analysed using Unisense SensorTrace Logger software. The sensor was calibrated using 2-point calibration with aerated tap water and an alkaline 0.1M ascorbate solution according to manufacturer’s recommendations. Analyses were performed in triplicate.

7. **RNA sequencing and analysis**

For RNA isolation, samples were transferred into separate tubes containing RNAlater®-ICE Frozen Tissue Transition Solution (Ambion, Carlsbad, CA, USA) and ground immediately using a pestle. Samples were stored on ice prior to RNA isolation. Three biological replicates were examined for each sample condition. Bacteria were grown until mid-exponential phase (OD_600_ of 0.2 to 0.3). Cultures were mixed with two volumes of RNAlater reagent and centrifuged at 8500 rpm for 10 min. The ground plant samples were centrifuged briefly at maximum speed. Total RNA was extracted using the Aurum^TM^ Total RNA Mini Kit (BioRad, USA) according to the manufacturer’s recommendations. Contaminating DNA was removed using the TURBO DNA-free^TM^ Kit (Ambion, USA). Additional RNA quality and integrity were checked by using the Pico 6000 RNA Kit (Agilent) and an Agilent 2100 Bioanalyzer instrument. Ribodepletion was done using the Ribo-Zero bacterial rRNA removal kit (Illumina), cDNA library preparation and sequencing on an Illumina HiSeq4000 system with a 75-bp paired-end read length were done at the Wellcome Trust Centre for Human Genetics (Oxford, UK). First strand cDNA synthesis incorporated dUTP. The cDNA was end-repaired, A-tailed and adapter-ligated. Prior to amplification, samples underwent uridine digestion. Libraries were prepared using Illumina TruSeq Stranded Total RNA Library Prep kit (Illumina, San Diego, CA, USA). Sequencing reads were trimmed using Trimmomatic [16] and mapped to the finished genome of *O. dioscoreae* LMG 29303^T^ using the EDGE-pro pipeline [17]. The resulting count matrix was analysed using the DESeq2 software [18]. Genes mapped by three or more reads and with Benjamini-Hochberg adjusted p-value < 0.05 and were considered differentially expressed (DE). Because the DEseq analysis yielded a very high proportion of differentially expressed genes (DEGs), additional analyses were performed using methods which are less prone to artefacts during library normalization. Specifically, we used Quantro [19] for quantile normalization of the dataset (p > 0.05), followed by qsmooth [20]. Final data matrices were analysed separately making use of both *limma-voom* and *limma-trend* packages [21]. Both *limma (voom/trend)* and DESeq2 pipelines are equally suitable and give similar results. Confirmation of differential regulation was done on genes of interest by RT-PCR (Figure S1). RNA-sequencing reads are deposited in the European Nucleotide Archive under study accession number PRJEB30089.

8. **Post-hoc validation of RNA-Seq experiments**

We verified the degree of genetic divergence between nodule bacteria and he cultured isolates by SNP analysis of the cDNA reads using Snippy with default settings (<https://github.com/tseemann/snippy>). Sampled nodule bacteria differed from strain LMG 29303^T^ by only 15 SNPs. However, we detected a cluster of 42 genes , which was present in strain LMG 29303^T^ but not in the nodule bacteria. This cluster of genes (ODI_R1825 – ODI_R1866) corresponds to a putative lysogenic bacteriophage, possibly acquired during or after isolation or passaging. The putative bacteriophage genes are only weakly expressed in strain LMG 29303^T^ (0.4% of all reads and may have a minimal effect on global gene expression patterns. In order to control for possible artefacts, all confirmatory RT-PCR experiments were conducted using both strain LMG 29303^T^ and strain R-67088, which has been shown by whole genome sequencing not differ from leaf nodule bacteria used in the experiment by only 6 SNPs and not to harbor the putative bacteriophage (data not shown).

9. **Quantitative Reverse Transcription-Polymerase Chain Reaction (RT-qPCR)**

First-strand cDNA synthesis was accomplished using the GoTaq® 2-Step RT-qPCR System (Promega, Madison, WI, USA), according to the manufacturer’s recommendations. Separate reactions containing no-RT and no-template were included as negative controls. qPCR was carried out on the LightCycler® 480 Real-Time PCR System (Roche Diagnostics) using gene-specific primers (Table 1). All oligonucleotides were designed using the CLC Genomics Main Workbench software. The primers were designed to be specific for *O. dioscoreae* genes (GenBank accession no. LT907988). The specificity of primers was tested by PCR, using gDNA of *O. dioscoreae* and *Dioscorea sansibarensis* nodules as the template in separate reactions. The total mRNA levels of genes of interest were normalized to those of the housekeeping gene *rpoD*, shown not to be differentially regulated in RNA-Seq experiments. All experiments were performed in triplicate.

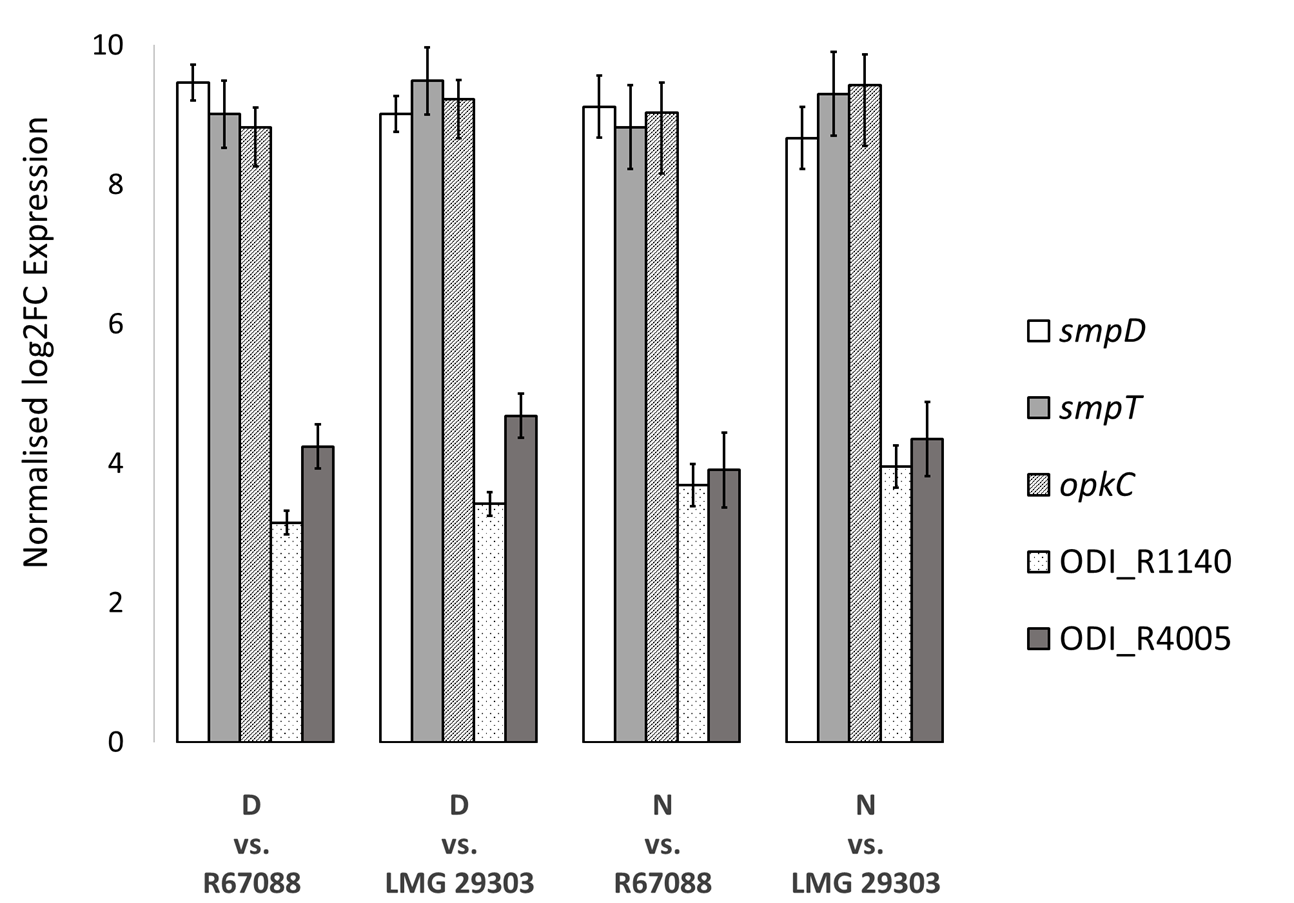

**Figure SI 1**: Validation of differential regulation of select genes by quantitative RT-PCR. D: leaf nodule RNA samples collected during the day. N: Leaf nodule RNA samples collected at night. Strain numbers represent cultures in AB medium supplemented with 0.2% sodium citrate and 0.05% yeast extract.

10. **Phylogenetic analyses**

A core genome phylogeny of all *O. dioscoreae* samples was constructed by aligning the protein sequences of all single-copy core genome orthologs, back-translating them to their nucleotide sequences, and concatenating them as described previously [22]. The resulting alignment, containing 3 555 045 positions divided over 3505 genes, was used to create a maximum-likelihood phylogenetic tree using RaxML v8.2.9 [23]. RaxML was run in rapid bootstrapping and best-scoring ML mode (-f a), performing 100 bootstrapping replicates, using the GTRGAMMA nucleotide substitution model. The position of the root was determined by repeating the analysis including *Achromobacter xylosoxidans* (NCBI accession nr. LSMI01000001) as an outgroup. Chloroplast genome alignments were constructed by mapping WGS reads to the draft genome of the *D. sansibarensis* chloroplast using REALPHY v 1.12 [24]. To avoid confusion due to sequencing errors in SNP-based phylogenies, samples for which estimated average coverage was below 20x were not included in the analysis, and the following REALPHY parameters were set: -readLength 150 –polyThreshold 0.9 to exclude ambiguous SNPs in repeat regions. The alignment generated had 125 175 positions, including 133 distinct patterns. The maximum likelihood model with the lowest Bayesian information criterion was selected using the model testing tool of CLC Main Workbench v.7.7.2. Chloroplast phylogenies were then reconstructed using the RAxML v8.2 program [23], using the GTRGAMMA substitution model, rapid bootstrap analysis and search for best scoring ML tree (-f a parameter setting) and 1000 bootstrap replicates. The same analysis was repeated by including the draft genome of the *D. elephantipes* chloroplast (NCBI RefSeq accession number NC_009601.1) to inform the root placement of the phylogenetic tree. Phylogenies of host and symbiont were visually compared and investigated using the TreeMap v. 3.0β program [25] by loading pruned species tree displaying only branches with support values > 70%. Reconciliation analysis was performed using the Jane v4 software [26] with default cost settings (co-speciation = 0, duplication = 1, duplication and host-switch = 2, loss = 1, failure to diverge = 1).

11. **Estimation of symbiosis age**

To determine lower bound for the age of the symbiosis, a dating analysis was performed to determine the approximate divergence time of our *Dioscorea* *sansibarensis* samples. Divergence time was estimated based on three plastid marker genes (*atpB*, *matK*, *rbcL*) and the *trnL* intron-*trnL* exon-*trnL*/*trnF* spacer, as described in Viruel *et al.* [27]. In short, Beast v.1.8.4. [28] was used with a Bayesian relaxed-clock approach, using the GTR+I+G substitution model, a Yule tree prior, and an uncorrelated lognormal molecular clock were used, allowing the rate of mutation to vary among partitions. Two MCMC chains were run for 100 million generations, sampling parameters every 10 000 generations. The same calibration points were used as in Viruel *et al.* [28], assigning lognormal prior distributions to fossil calibration points, and normal prior distributions for secondary calibration points.

Estimation of the age of divergence of the symbiont was performed by comparing the observed fixed mutation rate in the symbionts of one host lineage over a timespan of two years. This was combined with the estimated branch length of that branch in the core genome phylogeny to obtain an estimate of the divergence time. To increase representation of *O. dioscoreae* sequences, we included in our analysis whole genome sequences of strain LMG 29303^T^, isolated from a plant likely sampled in the Democratic Republic of Congo, although exact origin is uncertain, and strain R-67584 sampled from a plant of the live collection of the Botanic Garden Meise (accession CD-0-BR-1960001) also sampled in the DRC. Whole genome phylogeny of all genome sequences using RealPhy as described above placed the African isolates within clades of Malagasy genomes (Figure SI 2), and were not taken into account for divergence age calculation.

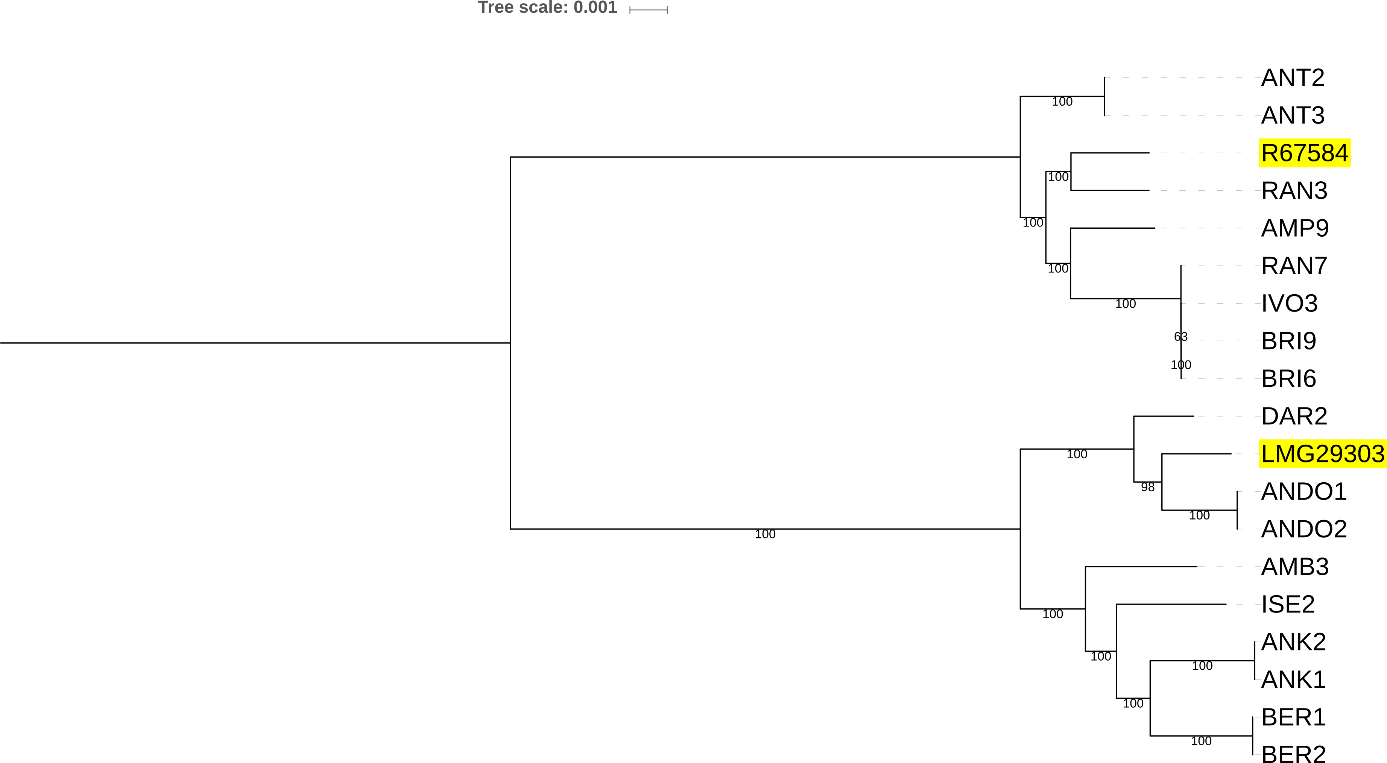

**Figure SI 2. SNP-based phylogeny of *O. dioscoreae* genomes.** The phylogeny was generated using the Realphy v1.12 with default alignment parameters and RAxML v8.2 with –GTRGAMMA and 100 bootstrap replicates (-f a -#100). Labels highlighted in yellow indicate stains isolated from plants collected in the DRC.

12. **Calculation of substitution rates**

Non-synonymous to synonymous substitution ratios (*d_N_*/*d_S_* or ω) ratios were calculated using the yn00 program of codeML, on the single copy core genome (1020 genes) of *O. dioscorea* (all strains) and several *Achromobacter* species (ASM23678v2, ASM21974v1, GCS2v1, ASM118959v1, ASM163968v1, ASM16583v1, Achr_xylo_C54_V2, ASM105105v1) *d_N_*/*d_S_* were averaged over all combinations within the same genus, discarding data with *d_S_* < 0.1 or > 2. To estimate the genome wide *d_N_*/*d_S_* of *O. dioscoreae*, this analysis was repeated a core genome of all *O. dioscoreae* genomes available consisting of 3563 genes. To identify genes under positive selection, site-specific *d_N_*/*d_S_* were calculated for all single-copy core genes of *O. dioscoreae* using *codeml.* Sites under positive selection were identified using the BEB (Bayes empirical Bayes) test comparing the M1a (neutral) and M2a (positive selection) models.

13. **Bacterial genetics**

To generate *O. dioscoreae* strain FID1, regions of homology flanking *smpD* (ODI_R1490) and a kanamycin resistance cassette from plasmid pKD4 [29] were amplified by PCR) using primers listed in Table S2. PCR amplicons were gel-purified using E-Gel Clonewell 0.8% SYBR Safe (Invitrogen) and assembled in an overlap PCR reaction. The final PCR fragment was cloned into a Gateway cloning vector *E. coli* pDONRpEX18 [30] using the BP Clonase™ II kit (Invitrogen, Carlsbad, CA, USA). The construct was transferred to *E.coli* Top 10 electro-competent cells for plasmid delivery using electroporation. The resulting plasmids were introduced to *O. dioscoreae* LF01, a spontaneous mutant resistant to nalidixic acid derived from strain LMG 29303^T^ by biparental mating using *E. coli* S17-1 as donor and transconjugants were selected by plating on TSA medium containing kanamycin (50μg/ml) and nalidixic acid (30μg/ml). Couter-selection of merodiploid clones was done by plating on medium containing 5% sucrose. Double cross-over events were verified by PCR.

14. **Chrome Azurol S (CAS) Assay**

Solid medium for the siderophore assay was prepared as described by Schwyn and Neilands [31]. Bacterial strains were grown as previously described, washed and suspended in sterile 0.4% NaCl to an OD_600_ of 0.01. Five µl from each of suspension of strains was spotted on CAS plates and incubated at 28°C for 48h. Halo formation was measured after incubation. Strains LMG 29303^T^, *O. dioscoreae* FID1, and LF01 all gave weak positive reactions, manifested as small, clear halos around the colonies . The *smpD* null mutant FID1 did not display significantly lower iron chelating activity than the parental strain, ruling out participation of the *smp* genes in siderophore production (data not shown).

15. **Growth on potassium galactonate as a sole carbon source**

Potassium D-galactonate was prepared by mixing a solution of 0.2M calcium galactonate (Sigma) and equimolar amounts of potassium oxalate (Sigma) in boiling water. After cooling, the calcium oxalate precipitate was removed by passing the solution through a 0.2 μm syringe filter. The solution was cooled and pH adjusted with potassium hydroxide prior to supplementation of culture medium.

Pre-cultures of *O. dioscoreae* strains LMG 29303^T^ and R-67584 were grown on TSA for 48h, washed in sterile 0.4% NaCl and resuspended in AB base medium without carbon source to an OD_590nm_ = 1. Fifty μL of cell suspensions were diluted in 5 mL of AB medium supplemented with 10 mM D-Galactonate or 10 mM sodium citrate and incubated at 28°C with shaking for 48h. Final OD_590nm_ of cultures grown on D-galactonate were 0.16 ±0.5 for strain LMG 29303T and 0.25 ±0.1 for strain R-67584. Final OD_590nm_ of cultures grown on sodium citrate were 0.13 ±0.2 for strain LMG 29303T and 0.175 ±0.2 for strain R-67584.

16. **Sequence analysis of the *smp* and *opk* gene clusters**

The central and largest gene of the operon, *smpD*, encodes a stand-alone module of a non-ribosomal peptide synthase (NRPS) (Figure SI 3). SmpD consists of a condensation domain, an adenylation domain and a phosphopantetheine-binding or thiolation domain, showing 29% identity to a module of bacitracin synthase 1 originated from *Bacillus licheniformis*. Substrate predictions of the adenylation domain are consistent with loading of a cysteine residue. Phylogenetic analysis and examination of the conserved catalytic residues place SmpD within the family of NRPS cyclization (Cy) domains (Figure SI4). SmpD may thus be involved in heterocyclization of a loaded cysteine to yield a thiazoline ring. The resulting heterocycle may be further oxidized to thiazole by the product of *smpF*, which encodes a putative flavin mononucleotide-dependent oxidoreductase. Unusually, SmpD lacks a thioesterase (TE) domain and release of the modified amino acid may occur through the action of a putative stand-alone TE encoded by *smpG*. Finally, SmpE is a hypothetical protein containing a heme oxygenase domain, possibly involved in redox tailoring reactions. At the 5’-end of the *smp1* operon, SmpA, SmpB and SmpC are likely also involved in tailoring reactions using acyl-CoA substrates.

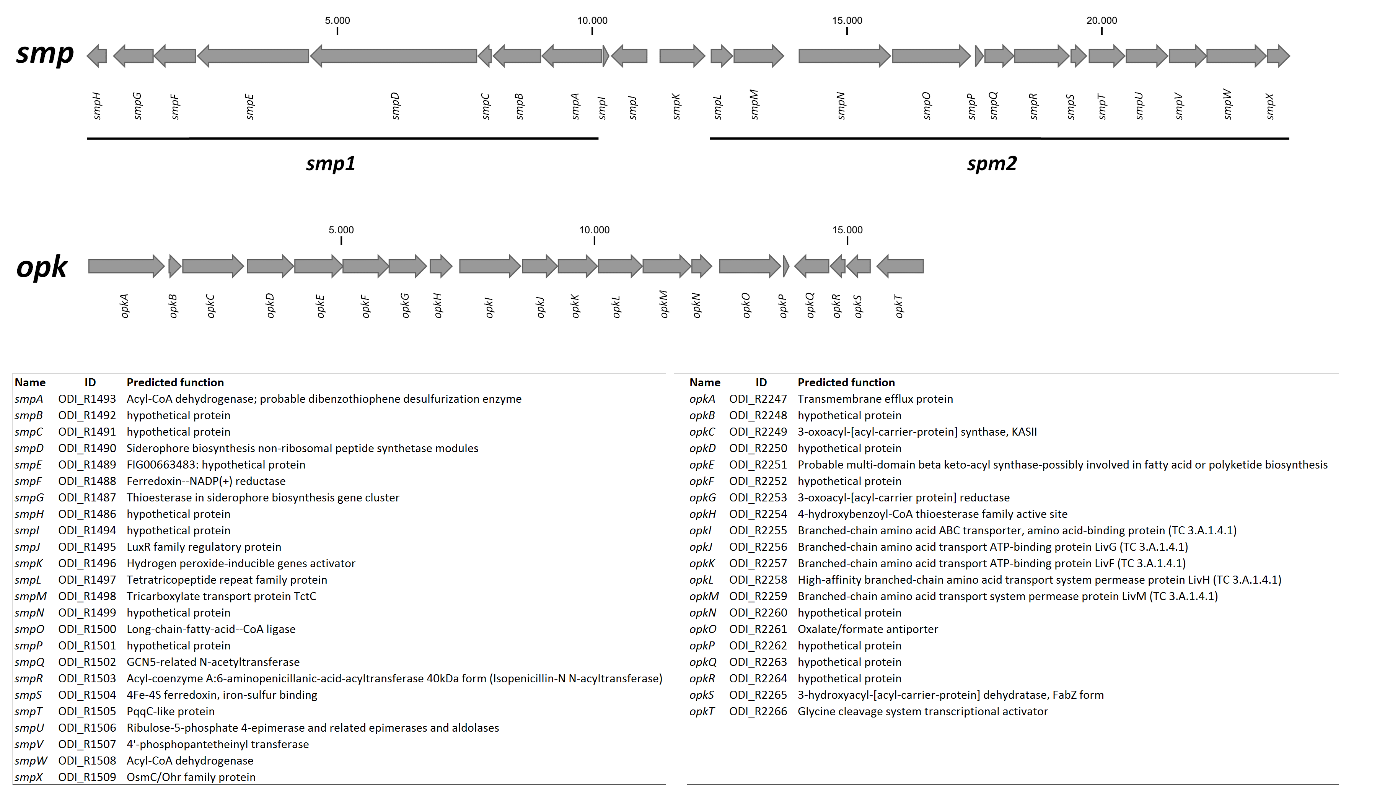

**Figure SI 3: *O. dioscoreae* gene clusters putatively involved in the synthesis of novel secondary metabolites.** Upper: Schematic overview of the *opk* and *smp* gene clusters, showing the unusual arrangement of biosynthetic operons (smp1 and smp2) surrounding the regulatory region. Table: Predicted function and locus tag identifier of the *smp* and *opk* genes.

The second operon of the cluster, *smp2*, is a highly unusual assemblage of redox enzymes and enzymes related to fatty acid biosynthesis. SmpL, SmpN and SmpP are hypothetical proteins with homologs of unknown function in *Burkholderiales*. SmpM shows homology (53.5% similarity) to the TctC periplasmic component of the Tct tricarboxylic acid transporter of *Comamonas testosteronii* but, homologs of TctA and TctB, the remaining components of the Tct ABC transporter are not encoded in the *smp* cluster or elsewhere in the vicinity. SmpO shows homology to long chain acyl-CoA ligase, and also has homologs in *Burkholderiales*. SmpR is a predicted peptidase of the C45 family, which includes acyl-CoA transferases. SmpV and SmpW are putative acyl-carrier protein and acyl-CoA dehydrogenase. SmpS and SmpT contain iron-sulfur binding and heme oxygenase domains, respectively, and are possibly involved in redox reactions. SmpU contains an aldolase domain, but also shows significant homology (35% identity) to NovR, an enzyme catalysing oxidative decarboxylation in the biosynthesis of novobiocin [32]. The biosynthetic product of *smp2* may thus be an acyl chain, which may possibly serve as substrate for the NRPS tailoring enzymes of *smp1*. The *smp1* and *smp2* operons are separated by two transcriptional regulators: a “solo” LuxR-family regulator containing an HTH and an acyl-homoserine lactone binding domain, possibly responding to an exogenous signal or endogenous acyl homoserine lactone quorum-sensing signal synthesized by the AHL-synthase of the LuxI/LuxR pair encoded just downstream of the *smp* cluster (ODI_R11512-1513). SmpK is a putative LysR family regulator.

The *opk* operon likely codes for enzymes linked to polyketide biosynthesis (Figure SI 3). OpkA is a putative transmembrane efflux protein from the Major Facilitator superfamily MFS-1 and shows high similarity (36% similarity) to a probable transporter of actinorhodin, a polyketide antibiotic produced by *Streptomyces coelicolor* [33]. OpkB and OpkQ are both putative acyl carrier proteins (ACP). A polyketide elongation module is comprised of two putative ketoacyl synthases OpkC and OpkD. OpkF, OpkG and OpkS are likely to be involved in further acyl chain modifications: OpkF is a putative AurF-family enzyme functioning as non-heme di-iron monooxygenase [34], OpkG is a putative 3-oxo-acyl-ACP reductase and OpkS is a putative 3-hydroxy-acyl-ACP dehydratase commonly found in polyketide biosynthetic clusters [35]. Finally, OpkH shows homology to a 4-hydroxybenzoyl-CoA thioesterase, possibly involved in releasing the polyketide chain from the acyl carrier. The *opk* cluster also codes for a highly upregulated putative ABC-transporter with homology to branched-chain amino acid transporters. OpkH shows limited homology (25% similarity) to the Leu/Ile/Val-binding protein of *Brucella melitensis*, but contains a well-conserved periplasmic binding protein (PBP) domain and a signal peptide, suggesting that the transporter is involved in uptake of a small molecule substrate rather than efflux.

**Figure SI4. Alignment of NRPS C and NRPS Cy domains.** Catalytic residues of NRPS C-domains are highlighted in green, and catalytic or conserved residues of NRPS Cy domains are highlighted in yellow. The protein sequence identifiers correspond to the identifiers in the NaPDoS database (<http://napdos.ucsd.edu/>, accessed November 2018).

```
Q9RCF7_VIBCH       ----------------MLLAQKPFWQRHLAYPHINLDTVAHSLRLTG---PLDTTLLLRA
TubCc              GGSLLAP-VARNGRLALSFAQQRLWFQEQLHPEAPANNLTGAVVFTG---PLHVAALLGA
epoBcy             -ESIVPAPAERHVPFPLTDIQGSYWLGRTGAFTVPS-GIHAYREYDCT--DLDVARLSRA
yersi1_C1_cyc      ----------------LTPVQHAYLTGRMPGQTLGGVGCHLYQEFEGH--CLTASQLEQA
yersi2_C1_cyc      ----------------LTPIQHAYWLGRTHLIGYGGVACHVLFEWDKRHDEFDLAILEKA
pyoch3_C1_cyc      ----------------LTPVQAAYVLGRQAAFDYGGNACQLYAEYDWP-ADTDPARLEAA
bleom9_C2_cyc      ----------------LTDVQRAYYVGREGGFALGGVSTHAYLEIEAP--RIDVARFTGA
pyoch2_C1_cyc      ----------------LSSVQQAYWLGRGAGEVLGNVSCHAFLEFRTR--DVDPQRLAAA
BACA_BACLIcy       LVTRAADPENIHEIFPLTGIQLAYLVGRDETFEIGGVATNLTVEFEA---DVDLNRFQLT
bleom9_C3_cyc      ----------------LTDIQRAYWLGRHRSLSLGGVATHTYLELDVE--DLDPGRLQTA
SmpD               PAHAAPPHAQAGPWFPLSAMQAAYLVGRGDTLALGRVSSHVYHEILME--GCDPDRLEAS
                                   :   *      .                            :  :

Q9RCF7_VIBCH       LHLTVSEIDLFRARFSAQGELYWHPF--SP--PIDYQDLSIHLEAEP-LAWRQIEQDLQ-
TubCc              VAALVRRHEALRTTLGEEGGVPYSLIGEPWQPALEVEALPGATVGERLEQAREVALAESR
epoBcy             FRKVVARHDMLRAHTLPDMMQVIEPK--VD-ADIEIIDLRGLDRSTREARLVSLRDAMSH
yersi1_C1_cyc      ITTLLQRHPMLHIAFRPDGQQVWLPQ--PYWNGVTVHDLRHNDAESRQAYLDALRQRLSH
yersi2_C1_cyc      WNQLIARHDMLRMVVDADGQQRILAT--TPEYHIPRDDLRALSPEEQRIALEKRRHELSY
pyoch3_C1_cyc      WNAMVERHPMLRAVIEDNAWQRVLPE--VPWQRLTVHACAGLDEAAFQAHLERVRERLDH
bleom9_C2_cyc      LRGVIARHPMLRAVIRPDGLQQVLTD--VPPYDVAVHDLRDLDEPARQRRRAALREEMSH
pyoch2_C1_cyc      AECVRQRHPMLRARFF-DGRQQILPT--PPLSCFDLQDWRTLQVDEAERDWQALRDWRAH
BACA_BACLIcy       LQKLIDRHPILRTIVFENGTQKILEA--TQRYTIETQDLRGFTEEEINVRILEQREKMTS
bleom9_C3_cyc      LRRLIDRHDALRLVVLPDGRQQILGD--VPPYLLAHTDLRGRADAE--AELARVREHMSH
SmpD               LCAVVARHEALRTIIDPEGRQAILPMQDVPTPLMTRHDHGRDDELAAQAAIHALRRRLSA
                         .   ::     :               .

Q9RCF7_VIBCH       RSSTLIDAPITSHQVYRLSHSEHLIYTRAHHIVLDGYGMMLFEQRLSQHYQSLLSGQTPT
TubCc              RRFALETEPHLRVRLLRLAEQQHVLVLSLHHIAADGVGLQVLEQELAALYGALSAGAEPR
epoBcy             RIYDTERPPLYHVVAVRLDEQQTRLVLSIDLINVDLGSLSIIFKDWLSFYE----DPETS
yersi1_C1_cyc      RLLRVEIGETFDFQLTLLPDNRHRLHVNIDLLIMDASSFTLFFDELNALLA----GESLP
yersi2_C1_cyc      RVLPADQWPLFELVVSEIDDCHYRLHMNLDLLQFDVQSFKVMMDDLAQVWR----GE--T
pyoch3_C1_cyc      ACAALDQWPVLRPELS-IGRDDCVLHCSVDFTLVDYASLQLLLGEWRRRYL----DPQWT
bleom9_C2_cyc      QVVPADLWPLFDVRVS-LGPTDALVHVGVDALICDAHSFGLVLAELAARYA----DPARR
pyoch2_C1_cyc      ECLAVERGQVFLLGLVRMPGGEDRLWLSLDLLAADVESLRLLLAELGVAYL----APERL
BACA_BACLIcy       KIIDPSVWPLFELKTFMLPGEKKYFFLNVDPLICDDSSMKRLIREFKQLYE----NPGLQ
bleom9_C3_cyc      EVRDASRWPLFDVRTHRLDDVRTRLHLSLDLLIADAHSVHVLTGDLLTFYA----DPDAA
SmpD               QVAPLARPCALEAVLVALPAGRHMLLVSHEGLHIDGLSMQILFADWAAAYA----RPDAT
                                    :      .    .    *  ..  .

Q9RCF7_VIBCH       AAF--KPYQSYLEEEAAYLTSHRYWQDKQFWQGYLREA----PDLTLTSATYDPQLSHA-
TubCc              LPPLPLQVADLADWQRRWVEGEEYQVQLAYWRRQLAGLTPL--EVPGDHPRPRIPSMRGA
epoBcy             LPVLELSYRDYVLALESRKKSEAHQRSMDYWKRRVAEL-PPPPMLPMKADPSTLREIRF-
yersi1_C1_cyc      AIDTRYDFRSYLLHQQKINQ-PLRDDARAYWLAKASTL-PPAPVLPLACEPATLREVRN-
yersi2_C1_cyc      LAPLAITFRDYVMAEQARRQTSAWHDAWDYWQEKLPQL-PLAPELPVVETPPE--TPHF-
pyoch3_C1_cyc      AEPLEATFRDYVGVEQRRRQSPAWQRDRDWWLARLDAL-PGRPDLPLRAQPDTR-STRF-
bleom9_C2_cyc      FPPLTADFRDHVLHQEALRGTAEYAAAERYWRERLPEL-PPGPELPLAVAPETLGTPRF-
pyoch2_C1_cyc      AEPPALHFADYLARRAAQR-AEAAARARDYWLERLPRL-PDAPALPLACAPESIRQPRT-
BACA_BACLIcy       LPSLEYSFRDYVLASINFKQTSRYQKDQQYWLDKLDHF-PSAPELPLKSDPAHVAKPSF-
bleom9_C3_cyc      LPPLGCSFRDYVLAVRAHAEGEPRRRALDHWRARLADL-PGPPGLPLRCRPEELTAPRF-
SmpD               LKALAPCVAPYVAAEQREREGPGWRQSRDRWLSRQARDGLHPPRLPLATDPDKLAQGLT-
                                                 *             :.

Q9RCF7_VIBCH       VSLSYTLNSQLNHLLLKLANANQIGWPDALVALCALYLESAEPDAPWLWLPFMNRWGSVA
TubCc              EVRAPLLSAPQAQVLRALGQ--GEGATLYMT-LLAALG------------VLLQRWTGQH
epoBcy             RHTEQWLPSDSWSRLKQRVG--ERGLTPTGV-ILAAFS------------EVIGRWSASP
yersi1_C1_cyc      TRRRMIVPATRWHAFSNRAG--EYGVTPTMA-LATCFS------------AVLARWGGLT
yersi2_C1_cyc      TTFKSTIGKTEWQAVKQRWQ--QQGVTPSAA-LLTLFA------------ATLERWSRTT
pyoch3_C1_cyc      RHFHARLDEAAWQALGARAG--EHGLSAAGV-ALAAFA------------ETIGRWSQAP
bleom9_C2_cyc      TRRSGRLDAASWTAVKDRAR--RAGLSPSGV-LLAAFA------------EVITAWSGRP
pyoch2_C1_cyc      RRLAFQLSAGESRRLERLAA--QHGVTLSSV-FGCAFA------------LVLARWSESA
BACA_BACLIcy       KKFSTFLDGHTWNELKKKAR--HHHLTPTSV-LCAAYA------------YILAYWSRQN
bleom9_C3_cyc      ARLTTGLGPDAWARLRRAAA--AAELTPAAL-ICAAFC------------DVLAQWSDTP
SmpD               ERFEASLDAQAWTRFQAHAG--AAGVTPAAA-VFAAYC------------DVLSRWDGTH
                         :       .           .                         :  *

Q9RCF7_VIBCH       ANVPGLMV-----------------NSLPLLRLFAQQ--TSLGNYLKQSGQAIRSLYLHG
TubCc              DMAVGSAAANRN--RPGLEGILGFLLNIVLLRLDLRGR-PRFRELLRQARRVCVEAYAHQ
epoBcy             RFTLNITLFNRLPVHPRVNDITGDFTSMVLLDIDTTRD-KSFEQRAKRIQEQLWEAMDHC
yersi1_C1_cyc      RLLLNITLFDRQPLHPAVGAMLADFTNILLLDTACD-G-DTVSNLARKNQLTFTEDWEHR
yersi2_C1_cyc      TFTLNLTFFNRQPIHPQINQLIGDFTSVTLVDFNFSAP-VTLQEQMQQTQQRLWQNMAHS
pyoch3_C1_cyc      AFCLNLTVLNRPPLHPQLAQVLGDFTALSLLAVDSRHG-DSFVERARRIGEQMFDDLDHP
bleom9_C2_cyc      RYSLMLTVFDRPPLHPDLGRIVGDFTSLSLLEVDHSRP-GDFTDRARALQRRLWQDLDHL
pyoch2_C1_cyc      EFLLNVPLFDRHADDPRIGEVIADFTTLLLLECRMQAG-VSFAEAVKSFQRNLHGAIDHA
BACA_BACLIcy       HFAINLTVFNRIPFHPDVKNMIGDFTSLMLLDIHAEENMSSFWRFALNVQDTLLEALEHR
bleom9_C3_cyc      RFTLNLTTFHRPALLPGVDDLVGDFTTTTLLGVDGE-G-DTFRDRARRLQDRIWEDLEHR
SmpD               AKTLNVTLAYRPPVSPDIDAAIGNFTRPVLATVTGTQP--RFDLRAKAAQAALMEALDLR
                                                *           .

Q9RCF7_VIBCH       RYRI----EQIEQDQGLNAEQ--SYFMSPFIN----------------ILPFESPHFADC
TubCc              ELPFEHLVEALQPGSE--RGDSSLYRVALAVSDTPWMPGHGLKLEGVQAQPLDFPR----
epoBcy             DVSGIEVQREAARVLGIQRGALFPVVLTSALNQQVVGVTS--------LQRLGTPVYT--
yersi1_C1_cyc      HWSGVELLRELKRQQRYP--HGAPVVFTSNLGRSLYSSRA--------ESPLGEPEWG--
yersi2_C1_cyc      EMNGVEVIRELGRLRGSQRQPLMPVVFTSMLGMTLEGMTI----DQAMSHLFGEPCYV--
pyoch3_C1_cyc      TFSGVDLLRELARRRGR-GADLMPVVFTSGIGSVQRLLGD--------GEAPRAPRYM--
bleom9_C2_cyc      AVGGVTVTRERALRHDARPGLLTPVVFTSDLPVGETAAED---ADGGEGWALGEPVYG--
pyoch2_C1_cyc      AFPALEVLREARRQGQP---RSAPVVFASNLGEEGFVPAA-------FRDAFGDLHDM--
BACA_BACLIcy       HYDGVDVIRNIAKKNGMNKKAVMPIVFTSVLSENPDDSFD-------SLVDFDNIHFF--
bleom9_C3_cyc      VVSGVEVLRMLRRERGTHDAVRMPVVFTSTLRAAGPAPRT-----APPAWRV-RPGYA--
SmpD               HFTALDAARHLASASPGQATLIVPYTFNFALADAPGHGP------LQGVETLGTHVHG--
                           .                 .   :

Q9RCF7_VIBCH       QTELKVLASGSAEGINFTFRGSPQHELCLDITADLASYPQSHWQSHCERFPR-FFEQLLA
TubCc              --------------------GVLDLDLHLWVYDT-GEGLTGRLEYAVDLYEEPTARRLLE
epoBcy             ------------------STQTPQLLLDHQLYEH-DGDLVLAWDIVDGVFPPDLLDDMLE
yersi1_C1_cyc      ------------------ISQTPQVWIDHLAFEH-HGEVWLQWDSNDALFPPALVETLFD
yersi2_C1_cyc      ------------------FTQTPQVWLDHQVMES-DGELMFSWYCMDNVLEPGAAEAMFN
pyoch3_C1_cyc      ------------------ISQTPQVWLDCQVTDQ-FGGLEIGWDVRLGLFPEGQAEAMFD
bleom9_C2_cyc      ------------------VSQTPQVHLDHQVAED-RGELVFNWDAVEDLFAPGALDAMFA
pyoch2_C1_cyc      ------------------LSQTPQVWLDHQLYRV-GDGILLAWDSVVGLFPEGLPETMFE
BACA_BACLIcy       ------------------STRTSQVYIDNQVYEI-NGGLYITWDYVEQIFEHEVIESMFD
bleom9_C3_cyc      ------------------ISQTPQVLLDHQVSES-DGRLVCTWDYVADAYPPGLIEAMFG
SmpD               ------------------VSQTPQVWLNLFVMRQ-RGGIVLQLDAVTGLFAPGVPACVAD
                                          :  :                              :

Q9RCF7_VIBCH       RFQQVEQDVARLLAEPAALAATTST----------RAIAS--------------------
TubCc              GFRQV---LEAVVEAPDRPVPELPV----LGEQERHQVLSGWNRTQRPYPREASVHGLFQ
epoBcy             AYVAF---LRRLTEEP--WSEQ--------------------------------------
yersi1_C1_cyc      AYCQL---INQLCDDESAWQKPFAD----MMPASQRA-----------------------
yersi2_C1_cyc      DYCAI---LQAVIAAPESLKTLASG----IAG----------------------------
pyoch3_C1_cyc      DFVGL---LRRLAQSPRAWTDGDAT-----------------------------------
bleom9_C2_cyc      AYTAS---LTRLARSPEAWRRPGTP----PLPTAQAA-----------------------
pyoch2_C1_cyc      AYVGL---LQRLCDST--WEQPADL----PLPWAQQAR----------------------
BACA_BACLIcy       QYIAV---IQKAVSGEDVSTIQMNEKSRQMISAYNDTDQS--------FDA-KPLHELFT
bleom9_C3_cyc      AFEAL---LASLAGHDDDAGHDDDA-----------------------------------
SmpD               SLGRL---LGRLADSPQAWQEAEFD----LLPATQRQARERANRTDKPVPV-ESLHAGFL
                           :
```
